# Integrating CSF and EV Proteomes Enhances Biomarker Discovery by Capturing Shared Variation Across Compartments

**DOI:** 10.64898/2026.09.30.754912

**Authors:** Bianca R. P. Brown, Xiaoting Li, Monica R Grasty, Andrew D Miranker, Gamze Gürsoy

## Abstract

Cerebrospinal fluid (CSF) contains brain derived proteins that can be leveraged as biomarkers for Alzheimer’s disease (AD) and other neurodegenerative diseases. However, converting protein intensities to a disease signal remains a challenge because physiology driven changes to the localization of proteins across CSF subcompartments are obscured when CSF is analyzed alone. Where subcompartment data, such as that from extracellular vesicles (EVs), are also measured, separate but parallel analyses of EVs and CSF can obscure their shared disease signal. Here, we present an integrated matrix factorization approach that combines proteomic signals from matched CSF and phosphatidylserine exposed EVs and nanoparticles (EV_ps_). In a ROS/MAP cohort of 48 participants, integrating CSF and EV_ps_ signals improved biological concordance across compartments and increased the number of candidate proteins distinguishing participants with AD from controls. Notably, the joint signal identified as many brain-related biomarker candidates as studies using substantially larger cohorts. By demonstrating that integration of CSF and EV_ps_ proteomes maximizes information from limited samples, this framework offers a path toward more efficient biomarker discovery across two compartments increasingly leveraged in neurodegenerative disease research.

## Introduction

Cerebrospinal fluid (CSF) is a complex biological matrix that contains brain-related proteins that can be leveraged as biomarkers for Alzheimer’s disease (AD) and other neurodegenerative diseases^1–7^. However, targeting and capturing these key proteins has posed a persistent challenge. The relatively low concentration of CSF, together with abundant background proteins, can make low abundance, disease-relevant proteins difficult to detect^8,9^. CSF provides a valuable window into brain biology, but its invasive collection means that clinical samples are often limited in both number and volume. It is therefore critical to maximize the information obtained from each sample; however, conventional bulk analyses can obscure disease-relevant signals arising from distinct molecular compartments within CSF. This challenge is further compounded by the imperfect nature of mass spectrometry–based proteomics, which preferentially captures high abundance proteins and underrepresents those at low abundance. Disease-associated changes can also be obscured by normal inter-individual variation, making it challenging to identify signals that consistently distinguish disease from biological heterogeneity. As a result, robust CSF biomarker discovery has often required large cohorts and sufficient sample volume, both of which can be limiting in clinical studies.

To overcome this signal-to-noise challenge, there has been increasing interest in broadening biomarker sources by collecting complementary data from partitions within the CSF. Extracellular vesicles (EVs) offer a compelling example. CSF EVs package and transport molecular cargo released from the central nervous system and are thought to reflect contributions from neurons, glial cells, vascular cells, and immune cells, including microglia and peripheral leukocytes^10^. Therefore, EVs have been better positioned than CSF by itself to report on neurodegenerative disease pathologies such as Alzheimer’s^11–13^ and Parkinson’s disease^14,15^. However, EV-based approaches face several technical challenges, including the low abundance of EVs in CSF, heterogeneity among vesicle populations, and incomplete characterization of their molecular composition and cellular origins^16^. A major limitation is the recovery of sufficient proteins from clinically reasonable volume^17,18^. These constraints mean that while EV-based biomarker discovery holds considerable promise, its broader application remains limited by current technical barriers.

To date, CSF and EV proteomes have largely been studied separately, despite capturing complementary aspects of the same underlying biology. Integrating these measurements may therefore reveal disease-associated signals that are obscured when each compartment is analyzed independently. As EVs arise from multiple origins, e.g. selective secretion by healthy cells, apoptosis by dying cells, with their cargo shielded from extracellular proteolysis, they represent a physiology curated view of the same fluid. For proteomics, these two fractions are not redundant. Whole CSF is biased toward reporting on the extracellular milieu while EVs report on the specific cells and cell states that created them (Fig. 1)^11,13^. Despite broad appreciation of this complementarity, CSF and CSF-derived EVs have invariably been studied in isolation. Even when both are collected from the same patients and used in the same study, they are analyzed in parallel^11,13^. This approach loses one of the most informative features of paired data collection: the covariance across compartments within individuals. A disease-associated abundance shift observed in both CSF and EVs carries different information when the two measurements co-vary across individuals than when the same marginal shifts occur independently.

**Fig. 1.**
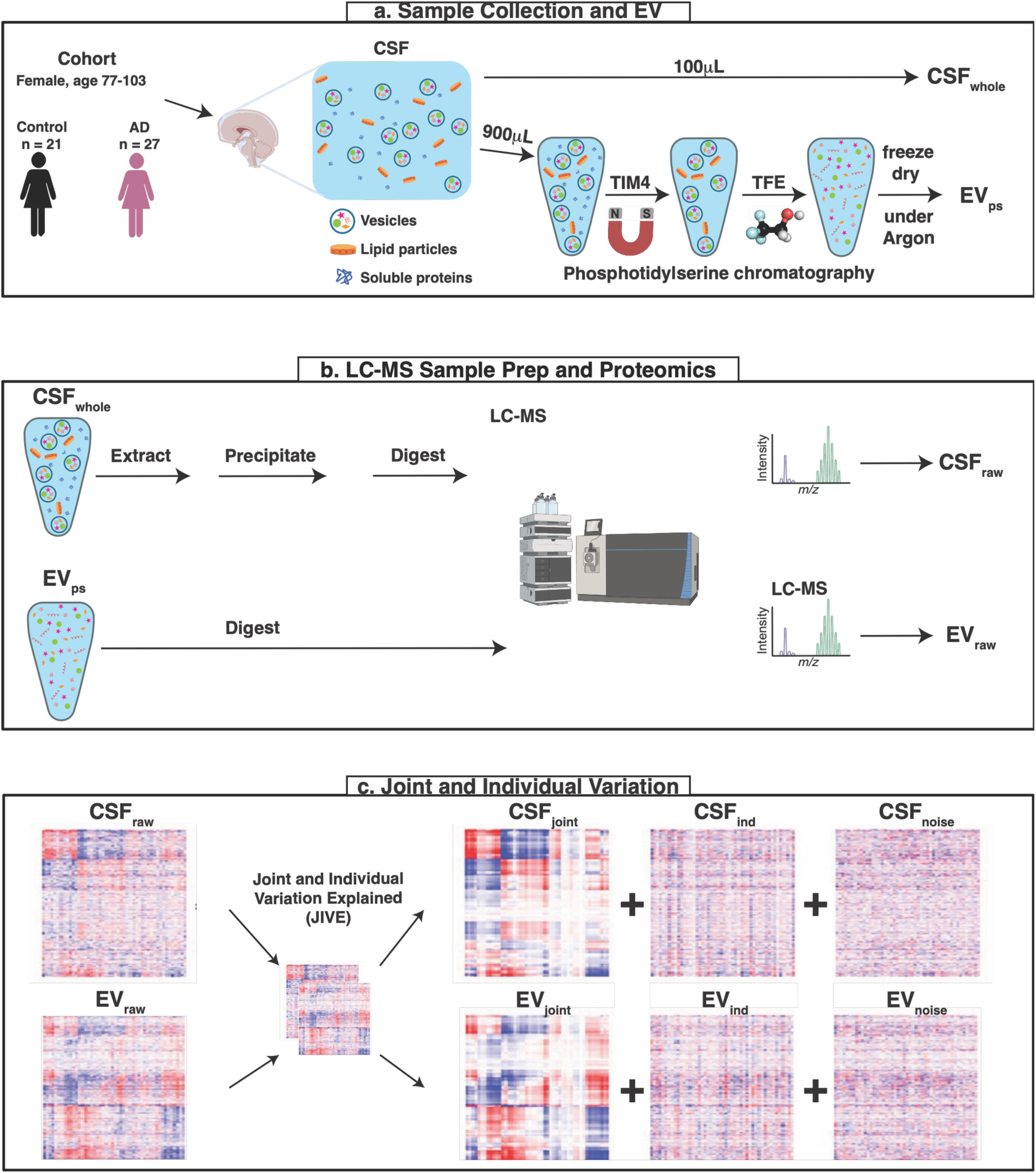
Schematic of the experimental and analytical workflow. (**a)** *Sample Collection and EV Extraction*: CSF is produced by the choroid plexus and circulates through the ventricular system and subarachnoid space, bathing the brain and spinal cord. CSF accumulates secreted soluble proteins, lipid particles, and EV secreted by choroid plexus epithelium and by cells throughout the central nervous system. CSF is collected from a cohort of patients. From each sample, an aliquot of CSF is exposed to TIM4-functionalized magnetic beads, yielding a phosphatidylserine enriched fraction (EVps). (**b)** *LC-MS Sample Prep and Proteomics*: CSF and EVps pairs are independently analyzed by label-free LC-MS proteomics, producing complementary measurements of the bulk fluid and its vesicular sub-fraction. (**c)** *Joint and Individual Variation Explained:* Paired data from the full cohort are co-decomposed using JIVE, which partitions each matrix into three components: a joint component capturing variation shared between CSF and EV, an individual component capturing variation specific to one matrix, and a residual (“noise”) component containing variation attributable to neither. The procedure yields six reconstructed datasets, three per matrix. Pairwise cross-correlations computed on the raw data are compared to those computed on the JIVE-denoised reconstruction, in which each matrix is rebuilt from its joint and individual partitions with the residual component excluded. As the linear algebra is not privy to relationships between proteins, covariation recovered and associated with known co-regulated complexes are used to validate the approach. JIVE partitioned data is then used to assess differential abundance between Alzheimer’s disease patients and controls.

We posit that capturing dispersed disease-associated signals across CSF-derived EVs and integrating them with molecular signals from CSF can improve biomarker detection. What is needed is a framework in which variation is partitioned rather than averaged, i.e. one that separates individual-level variation from covariation shared across partitions. Here, we establish such a framework and apply it to AD and cognitively normal participants from the Religious Orders Study and Memory and Aging Project (ROS/MAP) cohort^19,20^. For each subject we generated matched proteome measurements of whole CSF and CSF-derived EVs; the latter a non-trivial task given the small CSF volumes available from human donors, and from animal models more broadly. To maximize protein yield, we used a workflow we developed for non-human primate samples^21^ to enrich for the 10–30% of EVs with surface-exposed phosphatidylserine (PS), along with other PS-exposing nanoparticles^22^. This captures a reproducible, methodologically defined subcompartment of CSF (EVps).

To integrate paired CSF and EV_ps_ proteomics, we used Joint and Individual Variation Explained (JIVE), a matrix-factorization framework that separates molecular variation shared across sample compartments from variation specific to each compartment^23^. Originally developed for multi-omic data measured from matched samples, this structure is naturally suited to paired CSF and EV_ps_ analyses, where disease-associated signals may be distributed across both the soluble and vesicle-associated fractions. We, therefore, adapted JIVE as the statistical backbone of an integrated CSF and EV_ps_ biomarker discovery workflow, enabling us to distinguish convergent disease signals from compartment-specific molecular features. A practical implication of this work is that integrated CSF and EV_ps_ profiling may increase biomarker information of small cohort studies. We validate this premise by showing that the integrated framework better captures coordinated variation among biologically related proteins, suggesting that shared signals across CSF and EV_ps_ fractions reflect structured disease biology rather than technical noise. In a cohort of only 48 individuals, this approach recovered disease-associated proteins that substantially overlapped with biomarkers reported in studies with much larger cohorts. By integrating complementary protein information across CSF compartments, our framework promises to enhance the next generation of such studies, but also, can facilitate insight from small, deeply profiled disease cohorts by providing discovery power typically associated with consortium-scale studies.

## Results

We obtained CSF samples from 48 ROS/MAP participants, including 27 individuals diagnosed with AD and 21 cognitively normal controls (Table S1). EV_ps_ were prepared using a magnetic bead capture workflow that we previously optimized to minimize variability when dividing samples for orthogonal downstream analyses^21^. Our modifications are anchored in the use of a volatile, lipid-solubilizing, protein-disaggregating solvent: trifluoroethanol (TFE)^24^. TFE combines the normally separate steps of post-wash elution and EV lysis while eliminating the need for separate salt and detergent cleanup before liquid chromatography tandem mass spectrometry (LC-MS), thereby reducing sample handling and potential material loss in limited sample volume and low EV concentration samples used here.

EV_ps_ and their parent CSF samples were simultaneously processed for LC-MS, with volumes injected onto the column adjusted so that total protein from CSF and EV_ps_ samples was matched. We processed the raw mass spectrometry data using FragPipe^25^ (Table S2) and identified a total of 1,844 human proteins, including 332 proteins unique to the CSF dataset and 366 proteins unique to the EVps dataset (Fig. S1). We then used JIVE^23^ to integrate the paired CSF and EVps proteomes. JIVE identifies patterns of variation across samples that are coordinated between the two compartments while separating variation that is unique to either CSF or EVps. Importantly, the joint component does not require the same proteins to vary in the same way in both compartments; rather, each compartment can contribute different protein-level manifestations of the same underlying sample-level pattern. JIVE partitioned the preprocessed data (log2-transformed and per-protein mean-centered; CSF_raw_ and EV_raw_) into joint components representing this coordinated variation (CSF_joint_ and EV_joint_), individual components specific to each compartment (CSFind and EVind), and residual variation (CSF_noise_ and EV_noise_; Fig. 1, Table S3).

To distinguish the joint structure recovered by JIVE from that obtainable by chance, we performed a permutation analysis. We randomly reassigned sample IDs so as to disrupt only the matched pairing of CSF and EV_ps_ counterparts in the proteomic data (see Methods and Fig. S2). For correctly matched pairs, JIVE placed 37% and 24% of the variance present in CSF_raw_ and EV_raw_ into CSF_joint_ and EV_joint_, respectively (Fig. S2a). ID permutation wholly eliminated this. Across 100 independent ID randomizations, CSF_joint_ and EV_joint_ captured virtually none of the input variance (Fig. S2a-b). In marked contrast, the individual components (CSF_ind_ and EV_ind_) increased their capture of input variance from 38% and 23% in the original decomposition data set to 46% and 57% after permutation, respectively (Fig. S2a,c). Together, these results show that the joint variation recovered by JIVE depends on the true pairing of CSF and EVps samples, whereas disrupting this cross-compartment correspondence shifts the captured variance into compartment-specific components.

### Cross-correlation testing after JIVE resolves coordinated protein groups

Having established that the joint component depends on the matched pairing of CSF and EVps samples, we next asked whether this recovered structure organizes proteins into biologically coherent groups. If JIVE is capturing coordinated variation across compartments, proteins participating in related processes or molecular assemblies should exhibit stronger and more structured correlations in the joint components than in the corresponding raw data.

We therefore examined pairwise protein correlations within CSF_joint_ and EV_joint_ and compared them with those observed in CSF_raw_ and EV_raw_. Hierarchical clustering revealed multiple protein groups in the joint components that were not readily apparent in the corresponding raw datasets (Fig. S3). Because absolute protein abundance was not retained after preprocessing, proteins were ranked by their median values in the normalized and centered CSF and EV input matrices, and a representative subset was selected for correlation analysis.

The 50 highest-ranked proteins illustrate this effect (Fig. 2a,b; Table S4; Fig. S4). Three prominent clusters contained 13, 13, and 8 proteins, respectively, with within-group correlations approaching unity after JIVE decomposition. The relationships between clusters were also highly structured: groups *i* and *ii* and groups *i* and *iii* were anticorrelated, whereas groups *ii* and *iii* were positively correlated. By contrast, the corresponding proteins in CSF_raw_ showed substantially weaker intergroup relationships. Thus, JIVE did not simply increase pairwise correlation globally, but revealed a structured pattern of within- and between-group covariation that was largely obscured in the raw measurements.

**Fig. 2.**
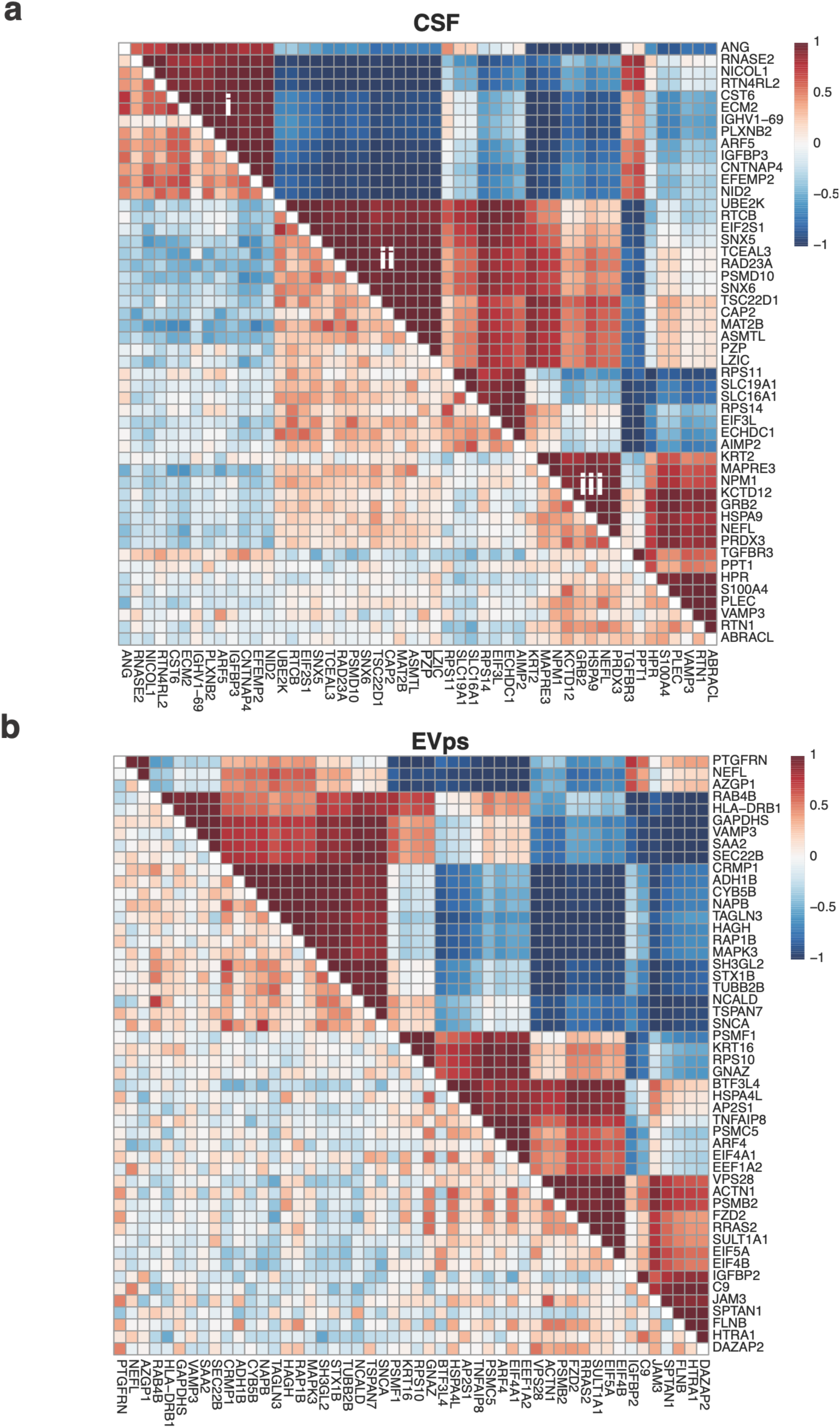
Cross-correlations reveal functional groupings in CSF and EV proteomes. Pairwise cross-correlation matrices of protein intensities are displayed as heat maps scaled from −1 (blue) to +1 (red). In each panel, the lower-left triangle reports correlations from the raw partition (CSFraw or EVraw), whereas the upper-right triangle reports correlations from the corresponding JIVE joint partition (CSFjoint or EVjoint), enabling direct visual comparison. Cross-correlations are shown among the top 50 proteins ranked by their median values in the normalized and centered CSF **(a)** and EVps **(b)** matrices. Coherent groups emerge in both CSF_joint_ and EV_joint_ that are not readily apparent in the raw data. Three groups, marked at right in panel a with Roman numerals (i–iii), are referenced in the main text. Briefly, **(i)** comprises 13 secreted or exported proteins, **(ii)** comprises 13 intracellular proteins spanning multiple tissue types and cellular functions, and **(iii)** comprises 8 intracellular proteins with greater CNS association.

Group *i* is dominated by proteins that reach the CSF through a signal-peptide-initiated secretory pathway and by regulated release from the plasma membrane. These include ECM proteins (NID2, EFEMP2, ECM2) and secreted factors (ANG, RNASE2, CST6, IGFBP3, NICOL1, IGHV1-69). Brain-enriched proteins in this group (PLXNB2, CNTNAP4) are receptors liberated by protease mediated shedding while RTN4RL2 is released in response to cleavage of its GPI anchor or by incorporation into an EV. ARF5 is a GTPase associated with membrane budding and a known constituent of plasma-derived EVs^26^.

Group *ii*, by contrast, comprises intracellular proteins with broad tissue distribution and dominated by protein-homeostasis machinery: the ubiquitin–proteasome system (the E2 conjugating enzyme UBE2K, the proteasome shuttle factor RAD23A, and the 19S-associated regulatory subunit PSMD10), the translation-initiation factor EIF2S1, and the tRNA-splicing/RNA ligase RTCB. Around this core sit proteins of endosomal sorting (the ESCPE-1 partners SNX5 and SNX6), transcription regulation (TCEAL3, TSC22D1), actin polymerization regulation (CAP2), and intermediary metabolism (MAT2B). This group also includes the less well-understood ASMTL, which does not traffic through the secretory pathway and is therefore more consistent with extracellular vesicle packaging or release through cellular turnover or damage. Lastly, one member, PZP, while clearly a part of homeostasis machinery, has anomalous localization. It is a secreted chaperone^27^ with reported Aβ binding activity, and its grouping with an otherwise intracellular cohort is unexpected.

Group *iii* is intracellular and spans several subcellular compartments and includes members more clearly associated with the CNS. This includes the axonal intermediate filaments NEFL, the brain-enriched microtubule tracking protein MAPRE3, and the GABA-B receptor subunit KCTD12. The remaining members are more broadly expressed and localize to cytosol (GRB2), nucleolus (NPM1), and mitochondrial matrix (HSPA9 and PRDX3). Lacking a route through the secretory pathway, their appearance in CSF implies release as part of the EV lumen or via cellular damage, the latter exemplified by NEFL, an established fluid marker of axonal injury. One member of group *iii*, KRT2, is an epidermal keratin and a known CSF contaminant. Here, it co-varies with the other group *iii* members, a pattern which we speculate is indicative of adsorption to lipid-bilayer particles, including EVs.

The changes to correlated intensity resulting from the application of JIVE, described above, can be compared to two limiting values. On the one hand, the average pairwise correlation of all proteins is 0.00±0.34 and 0.01±0.74 for CSF_raw_ and CSF_joint_ respectively. In contrast, pairwise correlations between subunits of known heterocomplexes are strong and become dramatically stronger after decomposition. The proteasome’s 14 core subunits yield cross correlations of 0.82±0.07 in CSF_raw_ and 0.52±0.19 in EV_raw_. After JIVE, these rise to 0.97±0.04 and 0.98±0.03 (Fig. S4). At much lower protein intensities, the 60S ribosomal subunit contributes 20 and 13 subunits to CSF_raw_ and EV_raw_, with average cross correlations of 0.55±0.12 and 0.17±0.20 (Fig. S4; Table S5); after joint analysis these improve to 0.89±0.12 for CSF_joint_ and 0.39±0.63 for EV_joint_. Notably, EV_raw_ data is required to enable the orthogonal analysis with CSF_raw_, with the higher correlations partition into CSF_joint_. Comparable gains hold for other complexes that can serve as positive controls including the TRiC chaperonin, the 40S ribosomal subunit, prefoldin, and the V-ATPase (Fig. S5; Table S5). In every case, inclusion of EV_raw_ followed by JIVE makes the assembled complex far more visible to cross correlation with the relationship best resolved in CSF_joint_. Clearly, JIVE is extracting coherent structure from the relationship between partitions, not from either one alone.

The use of obligate heterocomplexes as internal controls yielded an unexpected dividend in the form of illuminating outliers. Near-unity coherence was the expectation for the 60S ribosome, and it was met for most of the 20 subunits evident in CSF_joint_. Two deviated from the group and are interpretable (Fig. S4c). RPL11 was the dramatic standout, anticorrelated with the rest of the 60S. It has a well-characterized extra-ribosomal role in the ribosome-biogenesis stress response, binding MDM2 within the 5S ribonucleoprotein to stabilize p53, with downstream consequences for cell-cycle control. RPL7 is essentially uncorrelated with the other subunits, consistent with its moonlighting role as an extra-ribosomal translational repressor. RPL27A also deviates, but weakly and is not interpreted here. That the subunits departing most from complex coherence are those best-established outside the complex^28^ supports treating the correlation structure recovered by JIVE as reflective of underlying biology rather than statistical happenstance.

### Joint decomposition improves the number of AD-associated protein detected in differential abundance analyses

We next tested whether joint decomposition improves detection of disease-associated protein changes by performing differential abundance analysis on the original proteomic matrices (CSF_raw_ and EV_raw_) and their corresponding joint components (CSF_joint_ and EV_joint_). Because the joint components exhibit stronger dependence among proteins, we applied the more conservative Benjamini–Yekutieli (BY) procedure to CSF_joint_ and EV_joint_, while using the Benjamini–Hochberg (BH) procedure for CSF_raw_ and EV_raw_. Despite this more stringent multiple-testing correction for the joint datasets, JIVE substantially increased the number of AD-associated proteins detected: CSF_joint_ yielded nearly four-fold more significant proteins than CSF_raw_, while EV_joint_ yielded approximately 14-fold more than EV_raw_ (Fig. 3e,f).

**Fig. 3.**
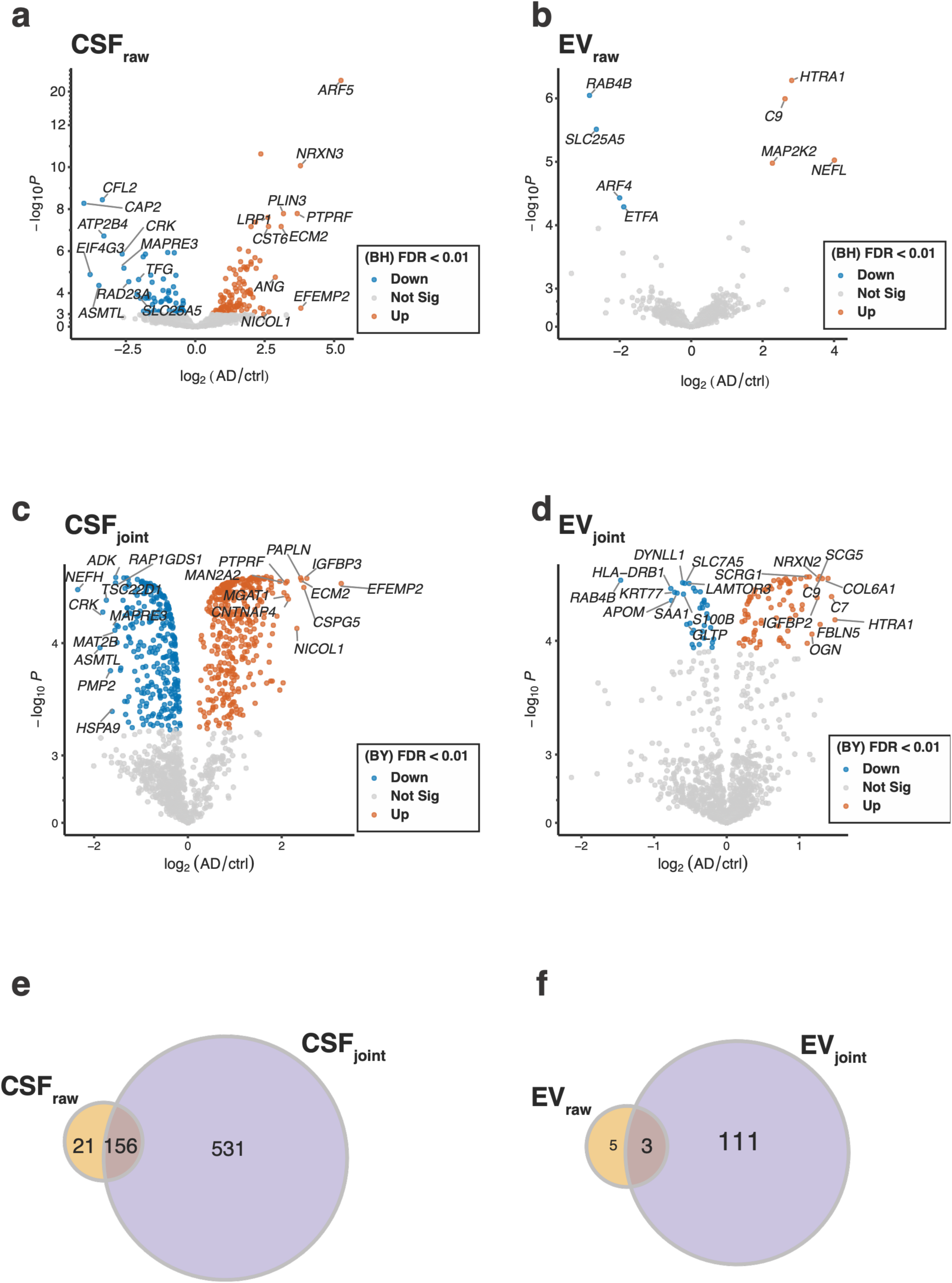
Differential abundance before and after JIVE decomposition. Volcano plots showing differential abundance results between AD and control individuals for **(a)** CSF_raw_, **(b)** EV_raw_, **(c)** CSF_joint_ and **(d)** EV_joint_. Proteins with statistically significant changes in intensity are shown in either orange (enriched in AD) or blue (enriched in controls). **(e,f)** Venn diagram of overlap of proteins identified as significant in CSF_raw_ vs CSF_joint_, and EV_raw_ vs EV_joint_ respectively, raw shown in yellow and joint in purple.

In CSF_joint_, 687 proteins were significantly differentially abundant between AD and control (Fig. 3; Table S6). The proteins most elevated in AD were EFEMP2, IGFBP3, PAPLN, ECM2, NICOL1, CNTNAP4, MGAT1, and MAN2A2, whereas those most elevated in controls were NEFH, ASMTL, CRK, TSC22D1, PMP2, HSPA9, MAT2B, RAP1GDS1, ADK, and MAPRE3. Several well-established AD-associated genes, including TREM2, BIN1, and PICALM, also reached significance. Notably, APOE protein abundance remained significantly associated with AD even after accounting for APOE genotype.

EV_joint_ yielded 114 significantly differentially abundant proteins, 80 enriched in AD and 34 in controls (Fig. 3; Table S6). The AD-enriched set included SCRG1, OGN, IGFBP2, SCG5, NRXN2, FBLN5, C9, COL6A1, C7, and HTRA1, implicating extracellular matrix remodeling, complement activation, and neuronal and secretory processes, with HTRA1 reflecting proteolytic and stress-related activity. The control-enriched set included RAB4B, HLA-DRB1, APOM, KRT77, SAA1, DYNLL1, S100B, SLC7A5, and GLTP. These are proteins associated with vesicle trafficking, immune surveillance, metabolic transport, and glial function, consistent with normal cellular homeostasis. These signals were confined to the joint channel: the individual components showed almost no AD/control differential abundance (Fig. S6; Table S6).

To assess the robustness of the differential abundance results, we performed an iterative leave-two-out resampling analysis. For each of the 27 × 21 possible combinations of removing one AD participant and one control participant, we repeated the differential abundance analysis. Median effect-size estimates (log fold change) across resampled datasets closely matched those obtained from the full cohort (Fig. S7), indicating that the observed differential abundance patterns were robust to the removal of individual samples.

Disease Ontology analysis indicates that the differentially abundant proteins surfaced by decomposition are associated with AD. The significantly differentially abundant CSF_joint_ proteins were associated with Alzheimer’s amyloidosis, cerebrovascular disease, brain infarction, and brain ischemia (Fig. 4a, Table S7), and the EV_joint_ proteins returned a dense neurodegenerative landscape including brain and cerebral infarction, retinal disease, and stroke (Fig. 4b, Table S7).

**Fig. 4.**
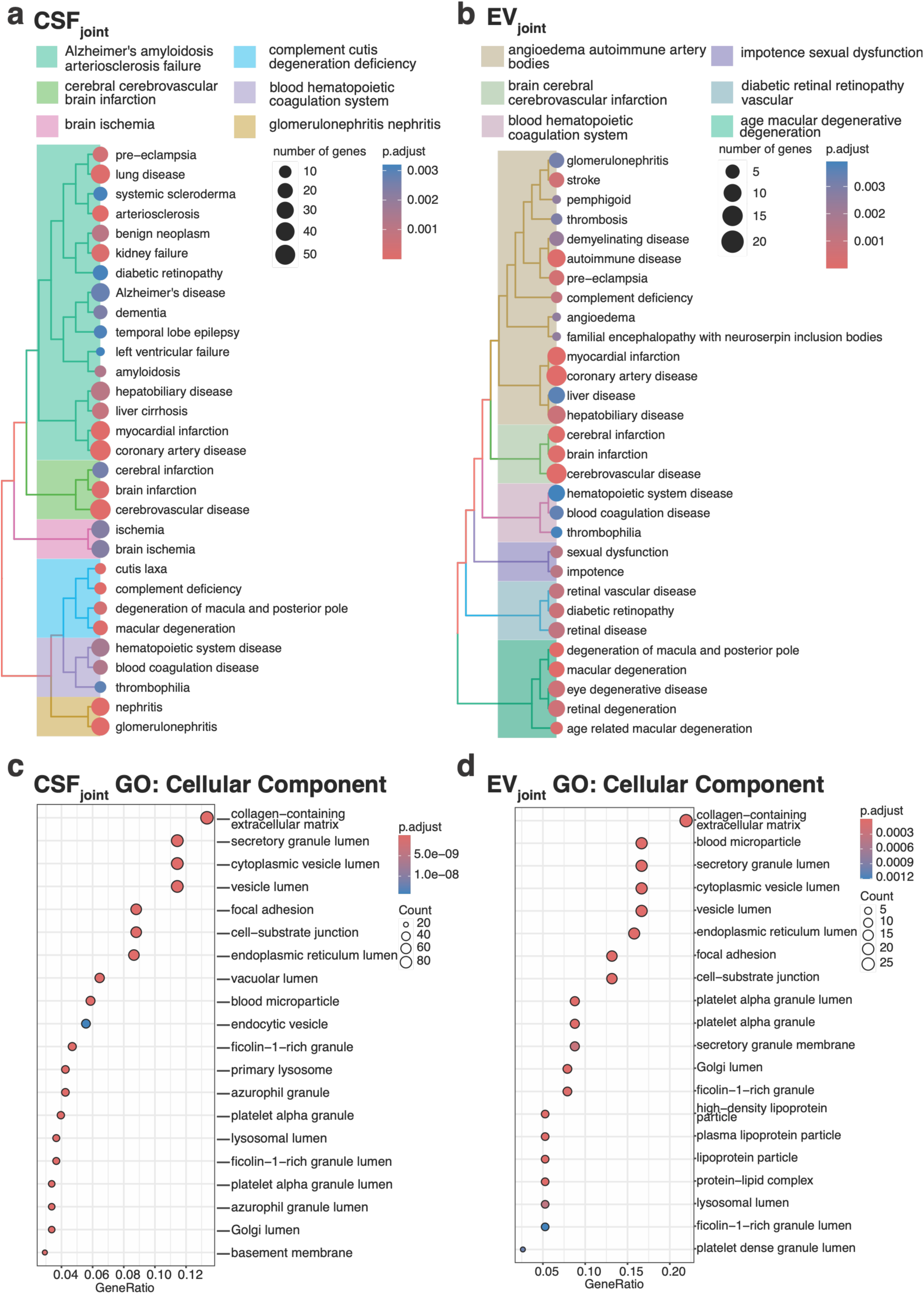
Disease and gene ontologies for AD-associated proteins in CSF_joint_ and EV_joint_. Tree plots show disease ontology terms for proteins significantly enriched in **(a)** CSF_joint_ and **(b)** EV_joint_ fractions. Top GO Cellular Component analysis is shown for **(c)** CSF_joint_ and **(d)** EV_joint_ fractions.

Corresponding analyses of the input matrices, CSF_raw_ and EV_raw_, returned no significant disease enrichment. The contrast is most informative for CSF_raw_ where 177 differentially abundant proteins returned no disease specific terms at all; the 21 proteins unique to CSF_raw_ were proteasome subunits (PSMA1, PSMA3, PSMA5, PSMB2, PSMB5), inflammation proteins (C1R, LTA4H), and the stress- and RNA-associated ST13 and HNRNPDL, but did not collectively implicate AD-specific biology. EV_raw_ returned only 8 differentially abundant proteins total, too few to support ontology-based enrichment testing, although the proteins did include ARF4, ETFA, MAP2K2, and NEFL. In contrast, proteins recovered after JIVE decomposition were enriched for amyloid, cerebrovascular, and ischemic processes, revealing disease-relevant biological structure that was not apparent in either input matrix.

The paired joint components also allowed us to compare disease-associated changes in the same proteins across CSF and EVps. Three cross-compartment patterns emerged. First, among proteins significantly differentially abundant in both components, CFL1, YWHAQ, and ALDOA showed opposite directions of association: each was more abundant in AD in EVps but less abundant in AD in CSF (Fig. S8).

Second, 262 proteins were present in both components but differentially abundant only in CSF_joint_ (Fig. S8, S9; Table S6). These included numerous vesicle- and trafficking-associated proteins (RAB5B, RAB6A, RAB6B, RAB7A, RAB8A, RAB10, RAB11B, RAB14, RAB21, RAB35, RALB, CDC42, VPS35, DNM1L, CLTA, CLTB, AP2A1, and LAMP2) and CNS-associated proteins that could plausibly be carried in CSF-derived EVs (NTRK2, NCAM1, NCAM2, CNTN1, CADM2, CADM3, CADM4, LSAMP, NEGR1, NTM, STXBP1, GPR37L1, CHGA, SCG2, SLC1A3, BCAN, CSPG5, PLP1, MBP, CNP, and TPPP3), together with a subset linked to lipid and lipoprotein biology (APOE, APOA4, RBP4, and NPC2). Many proteins shared between CSF and EVs therefore retain an AD-associated signal that is more strongly detected in CSF channel of the decomposition.

Third, 23 proteins were present in both components but differentially abundant only in EV_joint_ (Fig. S8, S9; Table S6). These included cytoskeletal and membrane-remodeling proteins (LASP1, FLNB, CAPG, and SH3BGRL); metabolic and redox-associated proteins (GLO1, GSTO1, GSS, TPI1, and HPRT1); NEDD8 and SUMO2, implicating altered protein regulation and post-translational modification; and extracellular or inflammatory proteins (KNG1, SAA1, APOM, CFH, ITIH4, and CP) that may show distinct disease-associated patterns when enriched in CSF-derived EVs rather than measured in CSF. Together, these patterns show that proteins common to both compartments can carry AD signals preferentially detected in one fraction over the other.

### CSF_joint_ identifies AD-associated CSF and brain proteins at a scale comparable to larger-cohort studies

We performed a cross-study concordance analysis to compare AD-associated proteins identified in CSF_joint_ with those reported in four published CSF proteomic studies with larger sample sizes: Bader et al. (2020; n = 60)^4^, Dammer et al. (2024; n = 300)^7^, Visser et al. (2022; n = 563)^5^, and Tijms et al. (2024; n = 606)^6^. We also compared our findings with an Alzheimer’s disease brain proteomic study, Markov et al. (2025)^29^ (Fig. 5; Table S8). Comparisons were restricted to proteins identified as differentially abundant between AD and control groups; for some studies, these analyses represented a subset of the total reported cohort (Table S8).

**Fig. 5.**
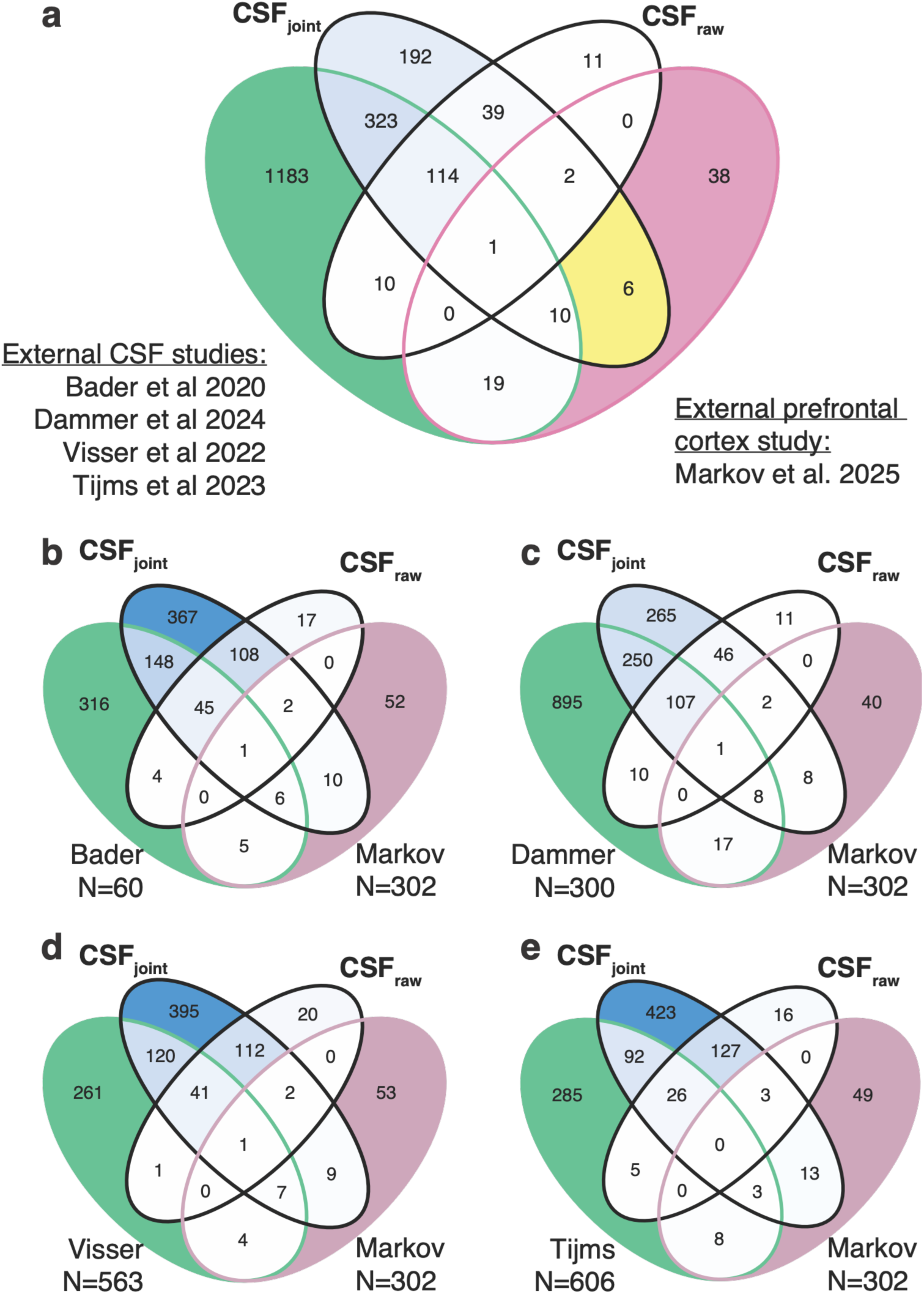
Comparison of proteins identified in this study with external datasets. (**a**) Combined comparison with Bader et al. (2020; *N* = 60), Dammer et al. (2024; *N* = 300), Visser et al. (2022; *N* = 563), and Tijms *et al* (2024; *N* = 606), illustrating the overall degree of overlap and the proportion of proteins unique to the present study. In yellow, six brain-associated biomarkers are uniquely identified in the CSF_joint_ analysis (HNRNPA3, CAPN1, NEFH, C1QBP, RAB5B, and AHNAK). Separate comparisons are shown for (**b**) Bader *et al*, **(c)** Dammer *et al*, (**d**) Visser *et al*, and (**e**) Tijms *et al*. Each CSF dataset was also compared with the brain (prefrontal cortex) proteomics findings of Markov *et al* (2025).

Joint decomposition substantially increased the number of AD-associated proteins corroborated by independent studies. Of the 687 proteins identified in CSF_joint_, 448 were reported in at least one of four external CSF cohorts, compared with 125 of the 177 proteins identified in CSF_raw_ (Fig. 5; Table S8). Although the proportional concordance was modestly lower for CSF_joint_ than CSF_raw_ (65% vs. 71%), joint decomposition more than tripled the absolute number of proteins supported by independent evidence. If the additional proteins identified by joint decomposition were primarily noise, they would be unlikely to show such widespread agreement with independent CSF proteomic studies.

The 239 proteins in CSF_joint_ absent from all four external studies, 198 of which are also absent from CSF_raw_, are also strongly informative. They include the AD-relevant NEFH, MBP, PLP1, CNP, LCN2, SPARC, and QPRT, together with a cohort of vesicle-trafficking and endolysosomal proteins: PICALM, VPS35, RAB7A, RAB10, RAB11B, RAB35, CLTA, and STXBP1. Functional enrichment analysis of these 239 proteins returns vesicle-associated and secretory compartment terms, alongside focal adhesion and cell–substrate junction (Fig. S10).

To place our own concordance in context, we assessed how well the external studies agree with one another. Agreement among them was uneven. Across the four comparator cohorts, 1,660 proteins were significant in at least one study; of these, 704 (42.4%) were significant in at least two, and only 29 (1.7%) were present in all four (Fig. S11). At the study level, 412 of 525 proteins identified by Bader et al. (78.5%), 354 of 419 identified by Tijms et al. (84.5%), 348 of 435 identified by Visser et al. (80.0%), and 597 of 1,288 identified by Dammer et al. (46.4%) were corroborated by at least one of the other three external studies (Fig. S11). Our 65% concordance observed for CSF_joint_ therefore falls within the range of concordance spanned by the external studies themselves.

Our CSF_joint_ identified between 10 and 16 AD related proteins from brain that were not present in each of the four external CSF studies. Similarly, each of the external CSF studies identified 5–17 proteins absent from CSF_joint_ (Fig. 5b-e). Compared against an AD prefrontal cortex proteomic dataset (Markov *et al*), our overlap with brain tissue was comparable to that of the external CSF studies: 19 of our significantly differentially abundant proteins were also identified in brain, eight of which were not reported in any of the four external CSF datasets (Fig. 5a). Of those 8, six only became evident after joint analysis (HNRNPA3, CAPN1, NEFH, C1QBP, RAB5B, and AHNAK). Plainly, orthogonal decomposition of our relatively small cohort is able to extract brain-derived signal efficiently and on a par with CSF-only studies with substantially higher patient enrollment.

To assess the sensitivity of our workflow on cohort size, we performed a stratified downsampling analysis and quantified the recovery of independently reported brain-associated AD biomarkers. Overlap increased steadily with sample size (Fig. S12): even a 30-sample subset recovered, on average, 5/19 brain-related biomarkers. Recovery of AD-associated brain proteomic signatures is still strongly sample-size dependent at 48 samples. The 19 proteins reported here should therefore be read as a lower bound of what is possible and a demonstration that biologically relevant brain-derived disease signal is accessible to small study enrollment cohorts.

## Discussion

The many sub-compartments of CSF, including but not limited to EV_ps_, hold valuable opportunities for investigating CNS disease biology and for biomarker recovery and discovery. Advances that make use of this clinically accessible biofluid have been hampered by the twin challenges of parallel acquisition of orthogonal data from single samples and subsequent data integration. We were therefore motivated to analyze paired CSF and EV_ps_ proteomes, treat them as a single dataset, and develop an analysis workflow scalable to an arbitrary number of sub-compartment fractions and mixed data types.

Our framework preserves matched sample structure by leveraging JIVE-based decomposition analysis. In doing so, we fundamentally increased the volume of information extractable from even modestly sized cohorts. Working with CSF/EV_ps_ as a pair carried an unexpected benefit: the set/subset relationship of CSF/EV_ps_ facilitated biological interpretation. Joint decomposition yielded a shared structure, expressed in each compartment as CSF_joint_ and EV_joint_, that revealed ontology supported AD-associations not readily apparent when either compartment was examined separately.

Four key findings anchor the above. First, JIVE assigned 37% (CSF) and 24% (EVps) of the variance in input proteomic intensities to the joint channels. Permuting the matched CSF–EVps pairings all but eliminated this assignment, establishing that the shared structure depends on genuine sample pairing. Second, decomposition enabled cross-protein correlations to resolve clusters not readily evident in the input data. For example, clusters whose members segregated by cellular origin and release route showed intra-cluster correlations near unity and inter-cluster correlations with characteristic sign and magnitude. Third, differential abundance testing identified 4-to 14-fold more significant proteins compared to the same statistical tests applied to the input data. Moreover, only the joint channels returned Alzheimer’s amyloidosis, cerebrovascular, and ischemic disease ontology terms. Fourth, external comparison indicated performance on par with far larger enrollment studies: 65% of our identified proteins were corroborated by at least one of four published CSF proteomes, which was comparable with how these studies compared with each other. In addition, 19 of our identified proteins overlapped with a published AD prefrontal cortex proteome, six of which became evident only after joint analysis. Taken together, these results establish the covariance of CSF and EV_ps_ proteomes as a measurable signal in its own right. Elements of disease biology obscured in either proteome alone become resolved once the pairing between them is preserved as an integral part of analysis.

The shared components, CSF_joint_ and EV_joint_, revealed biologically meaningful axes of variation. Using no prior information about protein localization, tissue origin, or physical interaction, the covariance structure separated a set of proteins into groups broadly consistent with soluble extracellular, intracellular, vesicle-linked, and brain-enriched biology. Internal heterocomplex controls behaved as expected, emerging as strongly coherent groups; notably, their deviations from coherence were themselves interpretable, as with the two ribosomal subunits highlighted above, RPL11 and RPL7, of well-established extra-ribosomal function. This calibration suggests that departures of correlation within expected groupings (be it by localization, shared control of expression, turnover or other mechanism), may mark underlying biology. Within this work, for example, outliers we cannot yet explain include PZP, an extracellular chaperone that clustered with intracellular proteins^27^. ARF5, by contrast, clustered with the secreted proteins despite its Golgi-associated localization, a pattern that is biologically plausible given its role in vesicle budding and its reported presence in plasma-derived EVs^26^. Such anomalous localization associations may of course prove fortuitous and false. Given how clearly the ribosome outliers map onto known biology, however, we suggest treating outlier behavior of the kind seen with ARF5 and PZP as a basis for prioritizing candidates for investigation rather than as noise to be discarded.

Integration of proteome data from CSF and its EV_ps_ subcompartment sharpened disease-relevant signals, recovering brain-associated AD proteins that conventional LC-MS CSF proteomics captures inconsistently. Benchmarked against four external studies, CSF_joint_ recovered more brain-associated AD proteins than three of the comparator datasets; only Dammer et al., using a patient cohort six-fold larger than ours, exceeded our total. In marked contrast, analysis of the present data without JIVE recovered fewer such proteins than any external study we examined. These gains therefore come from the decomposition of the combined data rather than from any property of this particular set of patient samples.

The brain proteins that CSF_joint_ recovered are predominantly intracellular, suggesting their appearance in CSF most plausibly reflects release or leakage following neuronal injury or cellular stress. Two are particularly instructive. NEFH and CAPN1 are unreported in the CSF datasets used for external comparison, yet targeted, affinity-, and activity-based platforms have reported both in the CSF of AD patients. This suggests that inclusion of EV_ps_ followed by joint decomposition confers a sensitivity approaching these specialized methods^30–32^. One feature of our NEFH result is unexplained: NEFH is elevated in controls rather than in AD, opposite to the direction reported by nonproteomic methods. This signal is internally coherent with co-assembling partners NEFL and NEFM positively cross-correlated (though neither meets our significance threshold on its own). Thus, while the sign difference is a consistent property of our data, its basis is unknown. The remaining unique proteins span the nucleus, mitochondria, and endosomal system, including HNRNPA3, C1QBP, RAB5B, and AHNAK^33–36^. Together they describe a diffuse signature of parenchymal compromise that falls below significance in CSF_raw_ but becomes resolved in CSF_joint_.

External overlap indicates that the proteins we identify as significant for AD in CSF_joint_ are not idiosyncratic. The four external studies corroborated 448 of our 687 reported proteins (65%), a concordance comparable to that achieved among the external studies themselves yet achieved by us with 48 patient samples. This supports the sensitivity of our workflow. The remaining 239 proteins, absent from all four comparators, have two non-exclusive explanations: some are proteins uniquely accessible through our combined CSF/EV_ps_/JIVE workflow, while others reflect postmortem release of intracellular proteins that antemortem comparator cohorts cannot capture. Non-overlap of this magnitude is in any case typical of biofluid proteomics, and we therefore put these 239 proteins forward as provisional biomarkers pending confirmation.

Beyond expanding the set of candidate markers, joint analysis resolves a distinct class of disease signal: sensitive to how a protein is partitioned between CSF and EV_ps_, rather than just a change in total abundance. Consider the case of a protein that lowers in total abundance in whole CSF of AD patients, but also shifts in its distribution from soluble to vesicle-associated pools. It would appear as a fall in one compartment and a rise in the other. CFL1, YWHAQ, and ALDOA approach this case: all three were differentially abundant in both CSF_joint_ and EV_joint_, but in opposite directions, rising in EV_joint_ among AD cases while falling in CSF_joint_. Such directional discordance argues against treating the two fractions as redundant measures of a single shared protein pool and indicates instead that AD variation distributes differently across CSF and EV_ps_.

Because CSF contains multiple EV subpopulations, not only EV_ps_, alongside soluble proteins and protein complexes, a protein detected in both CSF_joint_ and EV_joint_ may report on overlapping but non-identical molecular pools. Discordant AD-associated patterns could also arise if a protein is broadly EV-associated but preferentially carried by one EV subpopulation in AD and another in controls. Because we did not resolve multiple EV subpopulations here, these data cannot determine whether discordance reflects shifts in EV subtype abundance, altered cargo loading, or redistribution between vesicle-associated and non-vesicle-associated pools. Resolving that question is an opportunity in waiting, and one that expanding the set of subcompartments in the analysis would directly address.

Several experimental and analytical features of this study bound its present interpretation, while also pointing to new opportunities enabled by the combination of our experimental and analytic workflow. The ROS/MAP CSF analyzed here was postmortem ventricular fluid. We sought to lessen any potential impact of this by selecting samples for which postmortem interval was short (5.8 ± 1.8 hrs) and consistent between AD and control groups. This reduces the likelihood that postmortem effects can explain the observed disease-associated patterns^37^. Importantly, the total CSF volume used here, ∼1 mL, is ∼1/10^th^ the volume accessible by routine lumbar-puncture collections. In this study, EV enrichment was selective for a 10-30% subset of all EVs present. Parallel isolation methods might be used to isolate the remaining EVs, or even to further fractionate the EV pool by a physical or biochemical property. JIVE is not limited to only two orthogonal data sets, and such additional data sets can be readily incorporated.

The proteins uniquely identified by this study as AD-associated should be interpreted as discovery-level candidates requiring validation in prospectively collected cohorts with directed quantitation. In modest cohorts, the significance and even the direction of individual protein associations can prove sensitive to sample composition, particularly for proteins with small effect sizes. Our leave-two-out analysis, however, showed strong concordance between full-dataset logFC estimates and median resampled estimates, indicating that effect sizes were stable in this dataset. This sample-size constraint is in any case the very problem the framework was designed to address: maximizing recoverable biological information from limited clinical specimens. By leveraging matched CSF and EV_ps_ structure, JIVE nominates candidate signals that remain invisible when each compartment is analyzed separately.

As differential testing was performed using JIVE-reconstructed matrices, we asked whether decomposition might manufacture coherence or inflate significance rather than represent genuine biological structure. Several observations argue otherwise. Permutation of the correspondence between matched CSF and EVps samples collapsed the joint variation (Fig. S2), showing that the shared signal depends on true sample pairing, not on generic decomposition of two matrices. Moreover, we assessed the joint matrices using the Benjamini-Yekutieli procedure, which controls false discovery under correlated tests and therefore suits the dependence structure of low-rank components, while the raw matrices used the less conservative Benjamini–Hochberg procedure^38^. That the joint proteins remained significant under this more stringent correction further supports the robustness of the signal. Together, these results support interpreting the joint components as statistically derived but biologically informative summaries of shared CSF and EVps proteomic variation.

Joint decomposition of CSF and its PS-exposed subcompartment, EVps, yields a broader, systems-level account of AD pathophysiology than was accessible by either compartment on its own. The broader methodological contribution is that matched fractionation and joint decomposition can extract complementary proteomic information from a clinically accessible volume of CSF. In this study, a parent sample and just one of many potential matched partitions were sufficient to recover candidate biomarker signals comparable to those observed in much larger single-compartment studies. This suggests that limited-input, multi-compartment CSF profiling can improve the efficiency of biomarker discovery. More broadly, the framework provides a scalable route for integrating biologically defined CSF subfractions to support discovery across other diseases and/or injury of the central nervous system.

## Supplementary information

**Table S1 Metadata associated with samples from the Religious Orders Study and Memory and Aging Project (ROS/MAP) cohort.** Variables include sample ID, age, sex, genotype, disease status (control or Alzheimer’s disease), and post-mortem interval (pmi; hours after death) at sample collection. Cognition and additional clinical metadata were defined by physicians as part of the ROS/MAP study.

**Table S2 FragPipe output.** Protein-level quantification results generated using FragPipe for CSF and EVps. Each tab reports proteins with their associated unique peptide counts and intensities.

**Table S3 Joint and Individual Variation Explained (JIVE) output for CSF and EVps.** For each detected protein, the proportion of variance attributed to each JIVE component is listed: CSF_joint_, CSF_individual_, CSF_noise_, EV_joint,_ EV_individual_, and EV_noise_.

**Table S4 Correlation of the top 50 proteins**. Pairwise correlation for CSF_joint_, CSF_individual_, CSF_noise_, EV_joint,_ EV_individual_, and EV_noise_.

**Table S5 Correlation across biological complexes.** Pairwise correlation of a subset of biological complexes, including the TRiC chaperonin, 40S ribosome, 60S ribosome, proteasome core particle, proteasome cap, fibrinogen, prefoldin, tubulin, and V-ATPase.

**Table S6 Differential Abundance Analyses.** Protein level log fold change estimates between Alzheimer’s disease and control individuals for each JIVE component. Columns for the raw component include: logFC, AveExpr, t, P.Value, adj.P.Val, B, and Gene. The joint component includes the same columns with the addition of BY.adj.P to reflect the Benjamini-Yekutieli correction.

**Table S7 Functional enrichment analysis of significant differentially abundant proteins identified in the CSF_joint_, CSF_raw_, and EV_joint_ datasets.** Table includes Description, GeneRatio, BgRatio, RichFactor and FoldEnrichment, zScore, p value, adjusted p value and q value. geneID lists the proteins contributing to each enriched category, and Count indicates the number of significant proteins assigned to that term.

**Table S8 External study comparison.** The Table summarizes each external study including Study, Sample, Platform, Sample Size, Phenotype, Statistical approach and significance criteria, Proteins measured, and differentially expressed proteins identified. Per-protein overlap is also reported across Bader et al. (2020)^4^, Dammer et al. (2024)^7^, Visser et al. (2022)^5^, and Tijms et al. (2024)^6^, as well as an Alzheimer’s disease brain proteomic study, Markov et al. (2025)^29^, our CSF_joint_, our CSF_raw_, and the brain study.

## Supplementary Figures

**Fig. S1.**
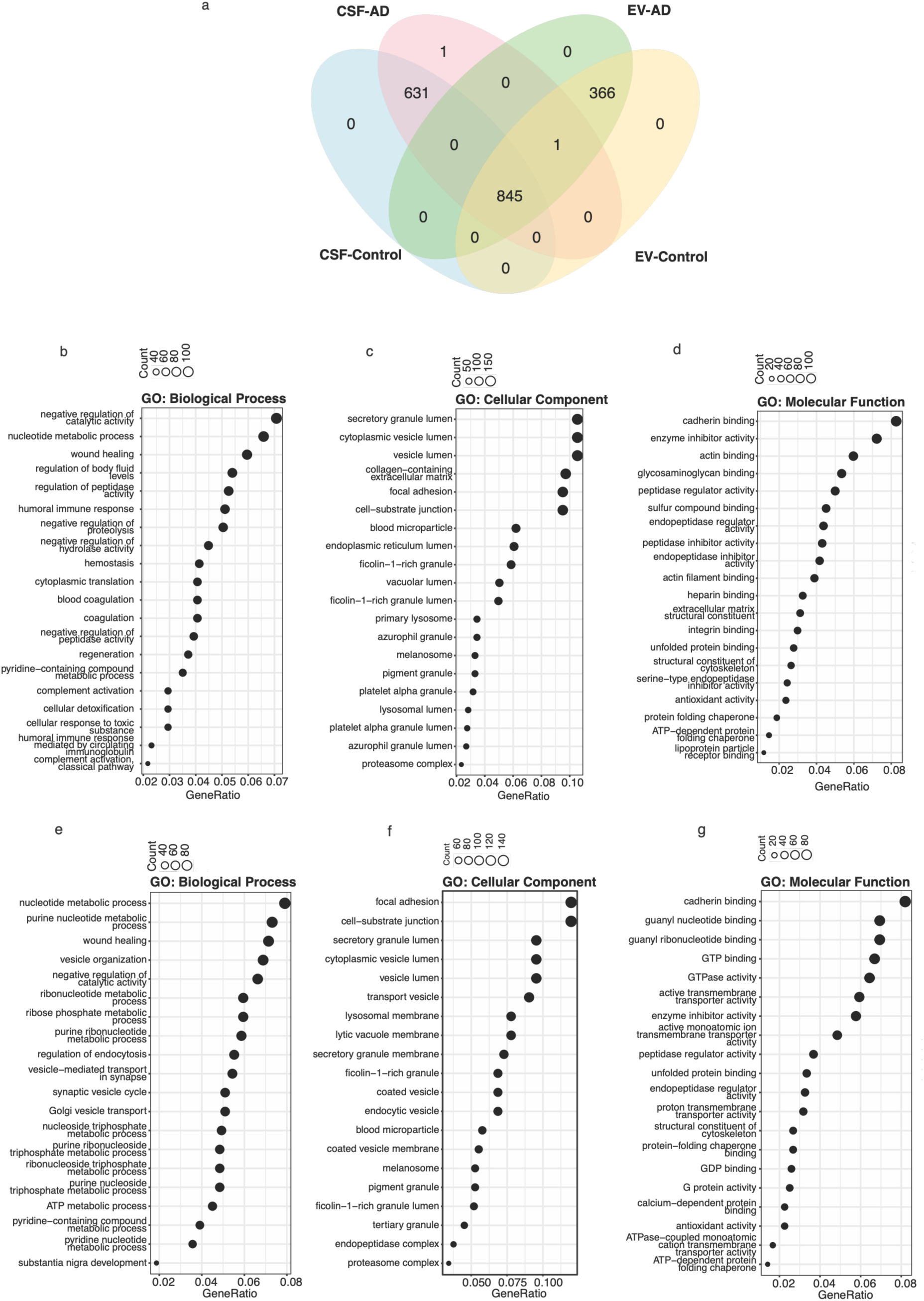
Functional characterization of proteins shared between CSF and EVps. **(a)** Venn diagram illustrating the overlap of proteins identified in CSF and EVps. Gene Ontology (GO) enrichment analysis of proteins identified in CSF_joint_ showing enriched **(b)** Biological Process, **(c)** Cellular Component, and **(d)** Molecular Function terms. GO enrichment analysis of proteins identified in EV_joint_ showing enriched **(e)** Biological Process, **(f)** Cellular Component, and **(g)** Molecular Function terms. All GO terms shown have adjusted p < 10^-8^.

**Fig. S2.**
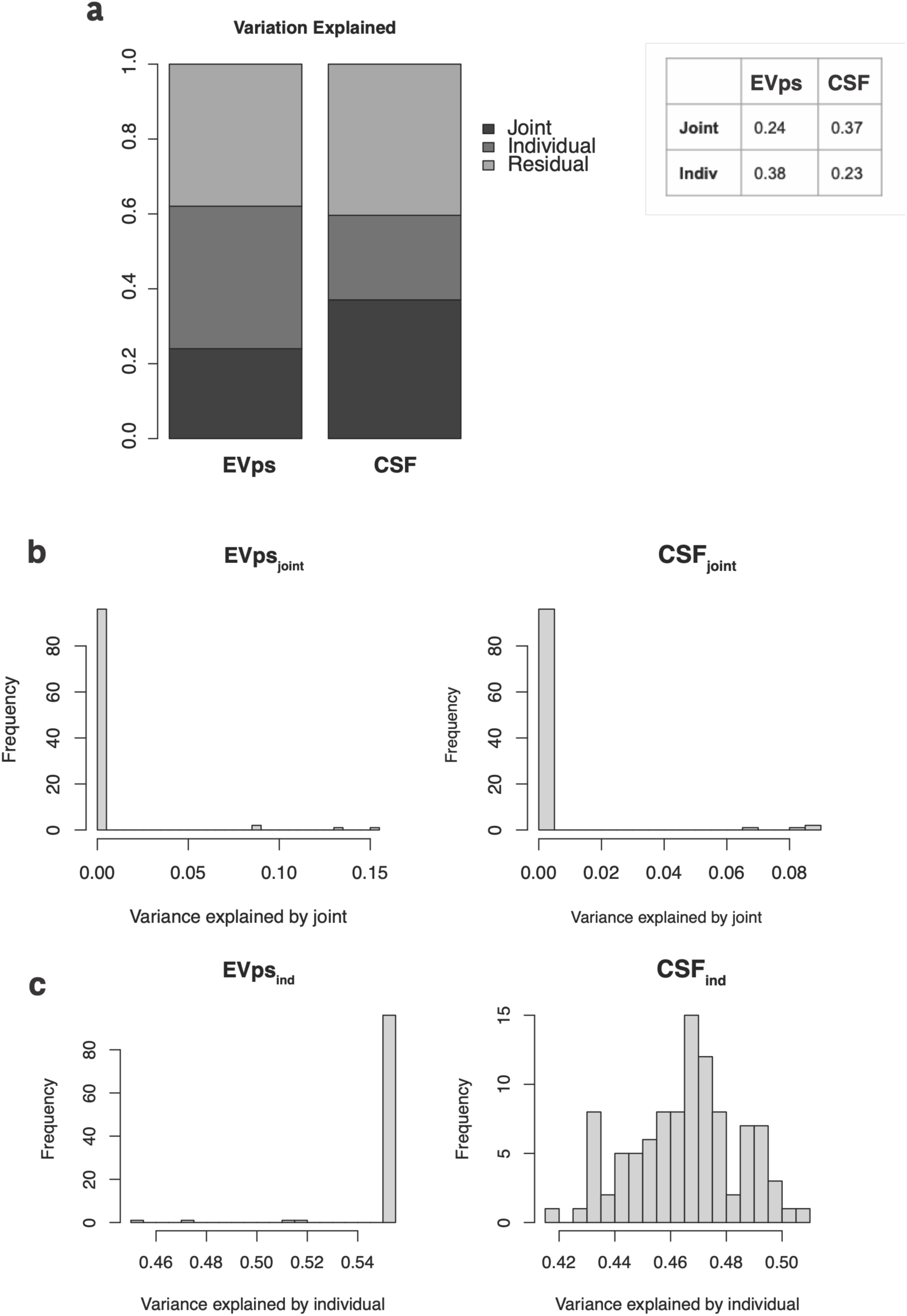
Robustness - permutation analysis. Sample columns of CSF matrix were randomly permuted for 100 times, to break correspondence between matched CSF and EV matched samples while preserving protein-level structure. JIVE was reapplied to each permuted dataset, and the variance explained by the joint and individual components was compared to the original decomposition data to assess the robustness stability of the shared joint structure. **(a)** Percentage of total variance explained by the joint, individual, and residual (noise) components for the original EV and CSF datasets. Exact percentages of total variance explained by the joint and individual components for the original JIVE decomposition in EV and CSF are shown to the table. **(b,c)** Distributions of the proportion of total variance explained across the 100 permuted datasets for the joint **(b)** and individual **(c)** components, shown separately for EV and CSF.

**Fig. S3.**
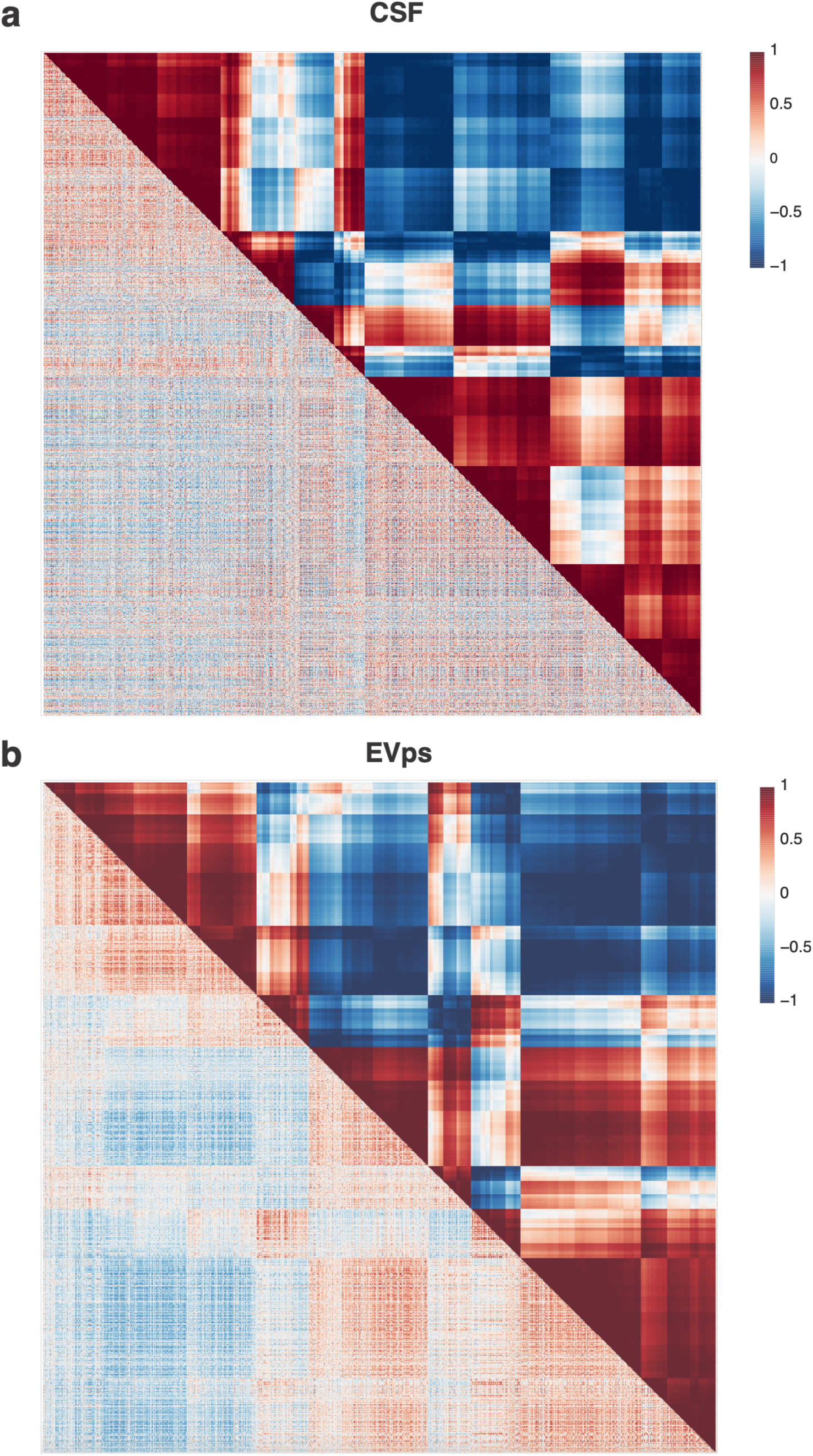
Pairwise protein correlations across the CSF and EVps proteomes. **a,b)** Pairwise correlations between all proteins measured in this work. In each panel, the lower-left triangle shows correlations calculated from CSF_raw_ or EV_raw._ Upper-right triangle shows correlations calculated from the corresponding JIVE joint component (CSF_joint_ or EV_joint_), for (**a**) CSF and (**b**) EV_ps_.

**Fig. S4.**
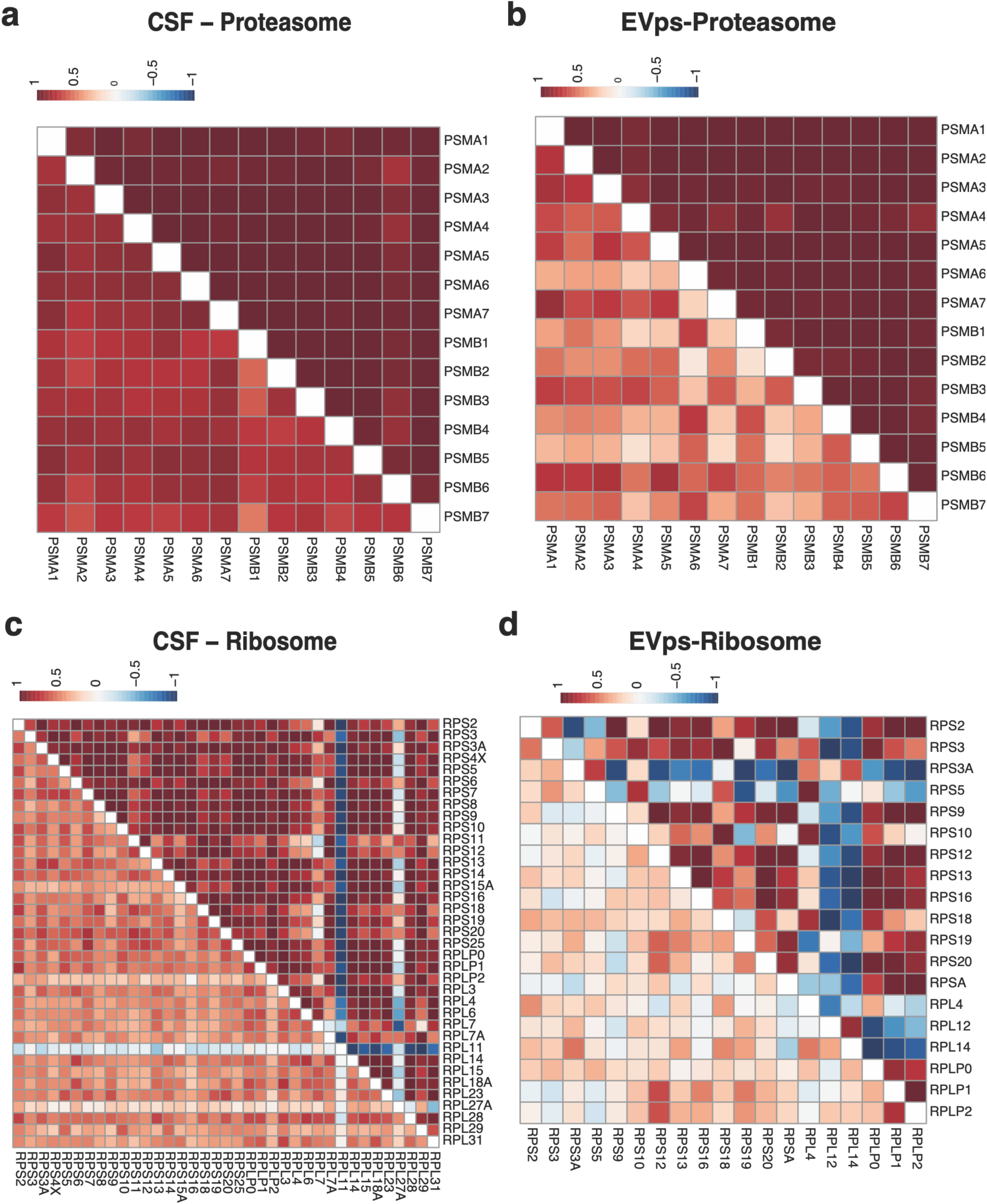
Correlation based on joint estimates and raw intensities for proteins that make up the proteasome and ribosome. **(a,b)** Cross-correlations among all 14 unique subunits of the 26S proteasome (**a,b**) and all measureable subunits of the 40S and 60S ribosomal particles **(c,d)** in CSF and EV_ps_ respectively. Joint analysis sharpens the within-complex correlation structure to near unity in both partitions demonstrating JIVE’s recovery of these known heteromeric complexes without prior input. *n.b.* Heatmaps were not clustered as they are obligate heterocomplexes.

**Fig. S5.**
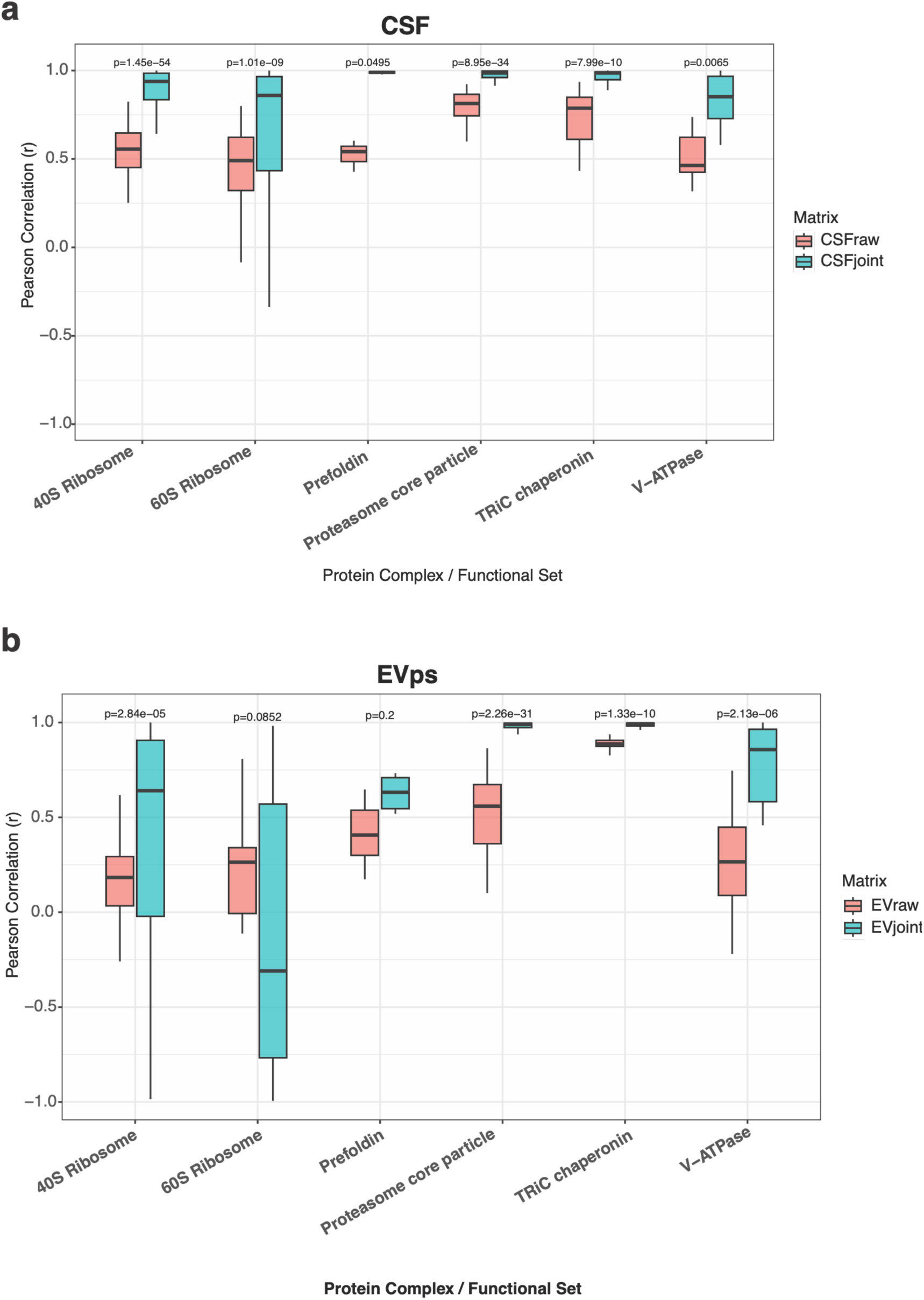
Correlation structure across protein clusters. Proteins comprising each cluster are listed in Table S5. For each cluster, we compared Pearson correlation values across the raw, joint, and individual components for **(a)** CSF and **(b)** EVs. Boxplots show the distribution of correlation values within each cluster, with Kruskal–Wallis p-values displayed above each panel to indicate whether correlations differed significantly across groups.

**Fig. S6.**
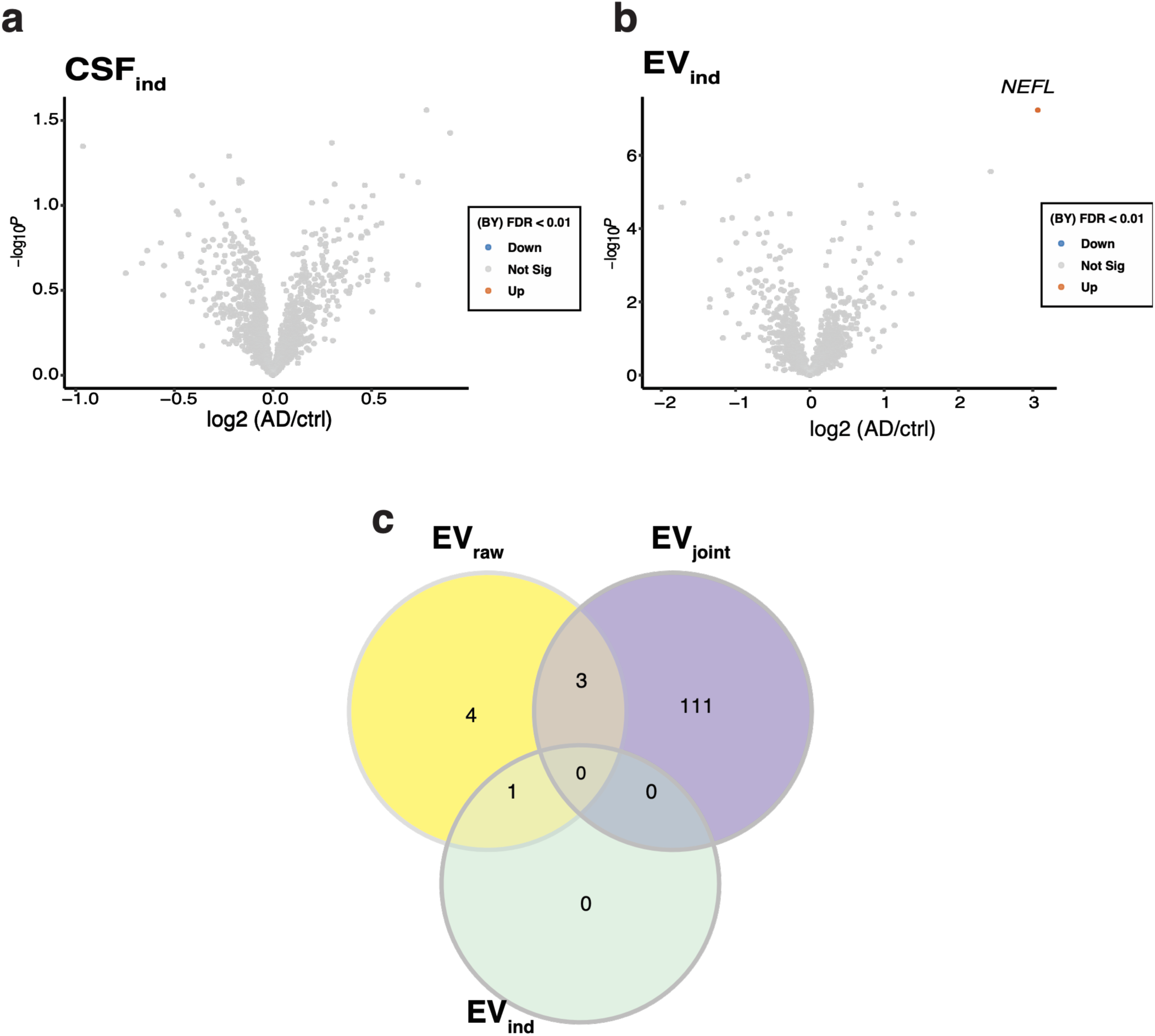
Differential protein abundance individual decompositions. **(a-b)** Volcano plots showing differential abundance results between AD and control patients CSF_ind_ and EV_ind_. Proteins with statistically significant changes in intensity are shown in either orange (enriched in AD) or blue (enriched in controls). **(c)** Venn diagram of overlap of proteins identified as significant in EV_raw_ vs EV_joint_ vs EV_ind_.

**Fig. S7.**
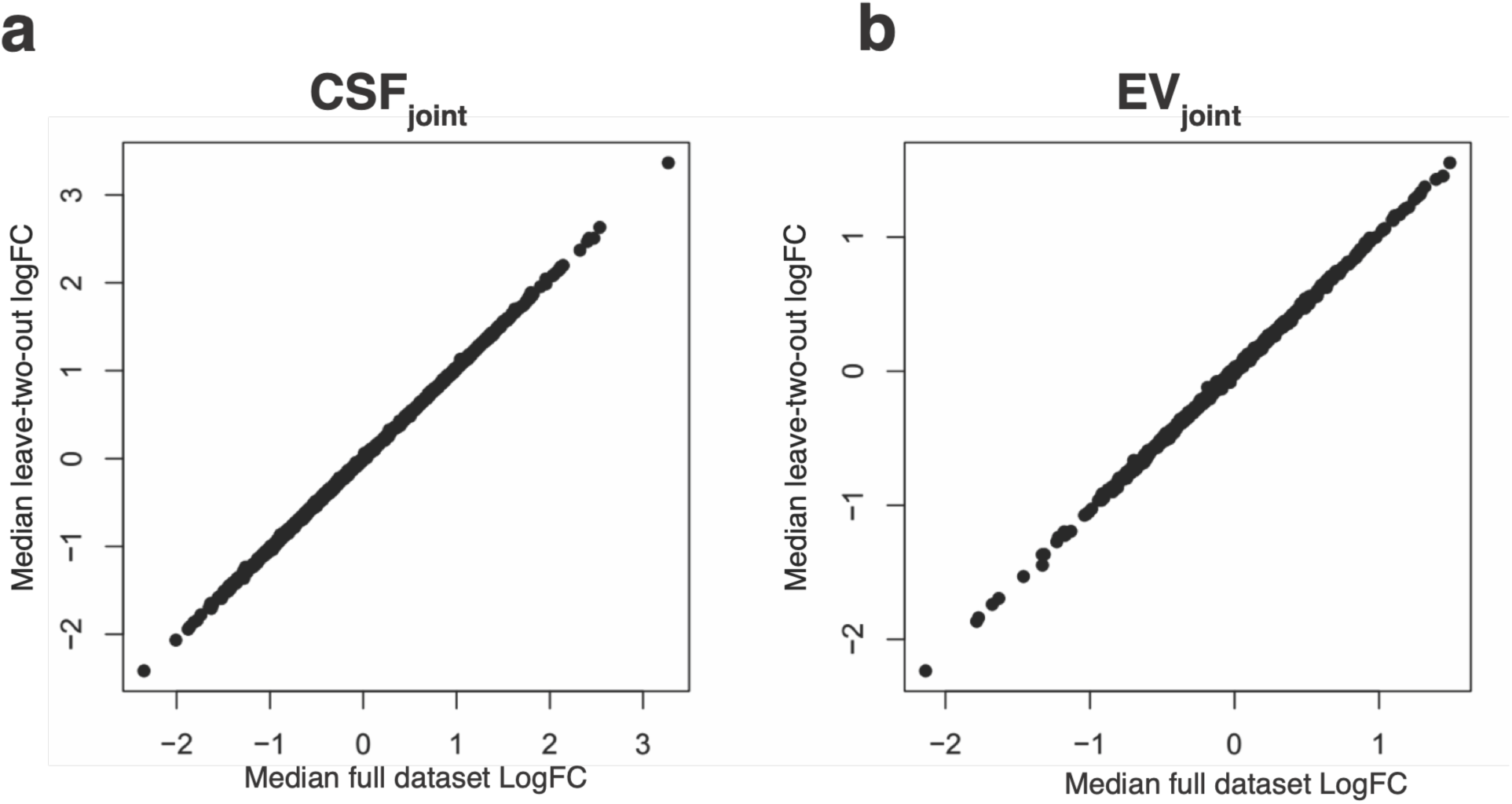
Leave-two-out cross-validation analysis of JIVE-derived differential abundance estimates. Scatter plot comparing protein log fold-change (logFC) estimates obtained from the full dataset (x-axis) with the median logFC across 567 leave-two-out cross-validation iterations (y-axis) for **(a)** CSF_joint_ and **(b)** EV_joint_. In each iteration, one AD and one control sample were removed, JIVE was re-applied to the reduced dataset, and differential abundance analysis was repeated using the same modeling framework as in the primary analysis. Each point represents a protein.

**Fig. S8.**
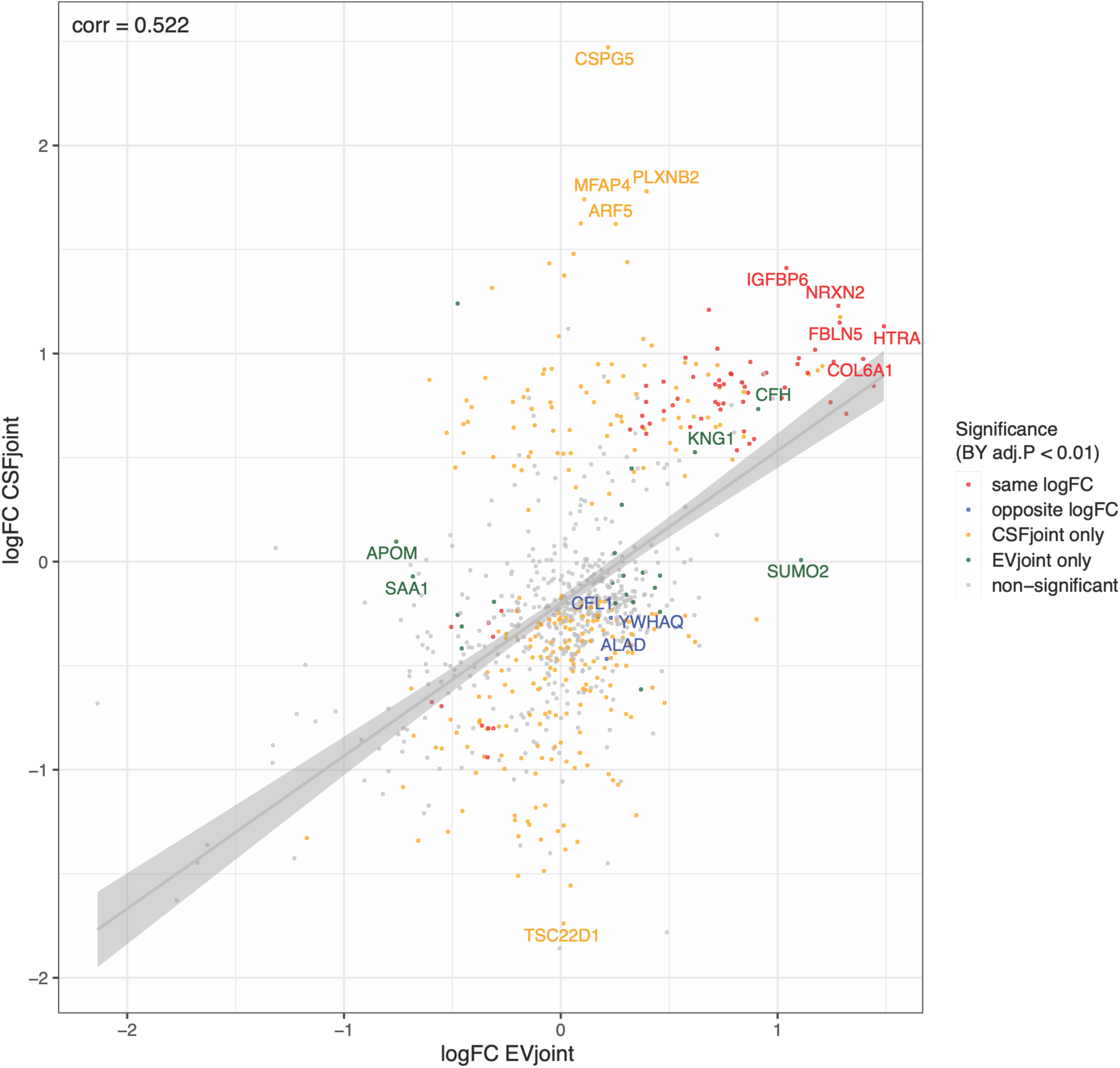
Correlation plot showing the log fold-change estimates for proteins detected in both CSF and EV_ps_ based on differential abundance analyses. Plot highlights the relationship between compartment-specific effect sizes. Red are proteins that had the same log fold change for CSF_joint_ and EV_joint_ and blue are those with opposite log fold change. Yellow represents those that are only significant in CSF_joint_ and green EV_ps_ only. Gray are the proteins that are not significant.

**Fig. S9.**
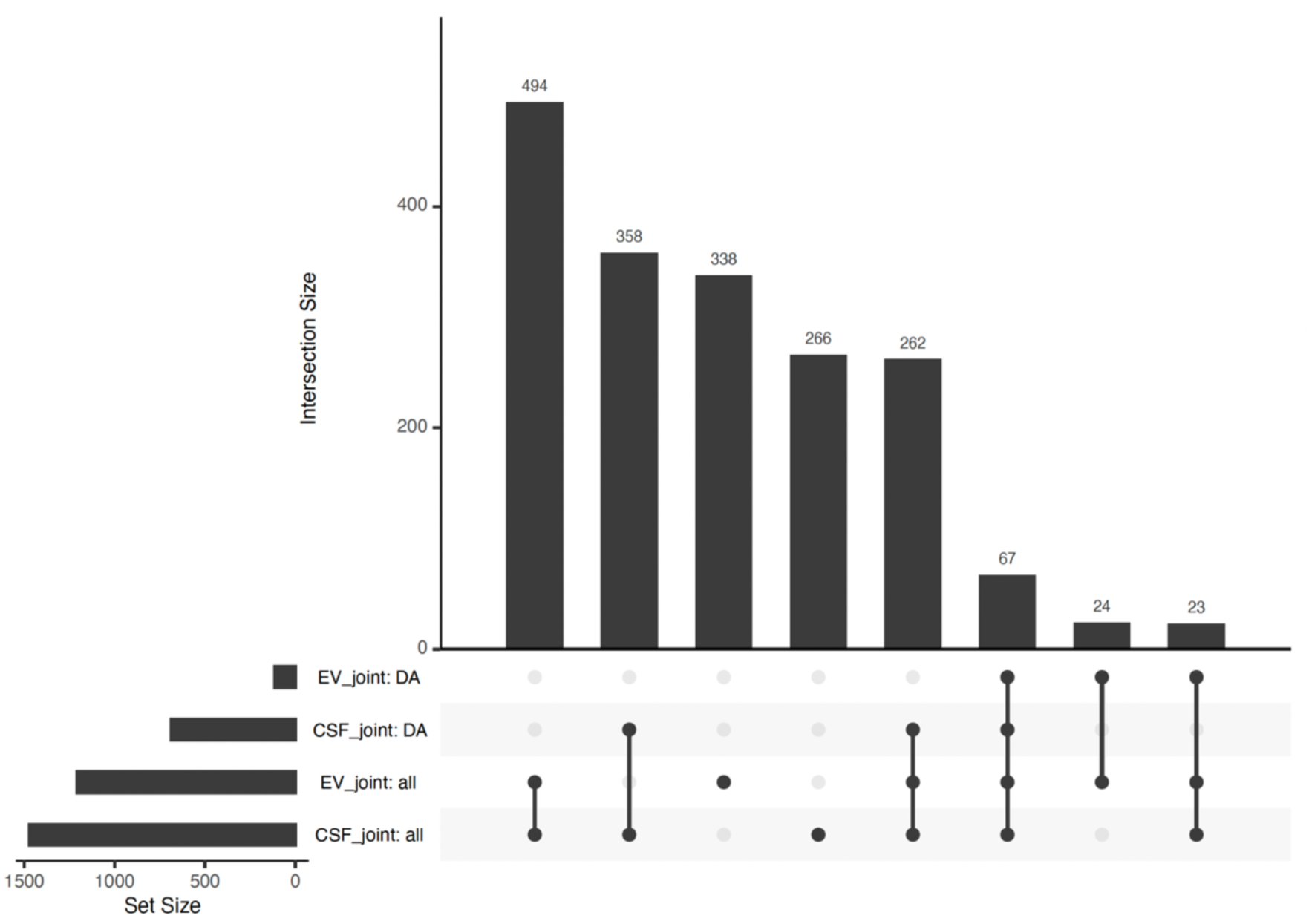
Overlap of proteins detected in both CSF_joint_ and EV_joint_. Upset plot showing overlap to identify proteins that differ in their differential abundance patterns across CSF_joint_ and EV_joint_. Proteins included in both joint datasets may be significantly differentially abundant in CSF_joint_, EVj_oint_, or both, highlighting proteins with shared detection but compartment-specific disease-associated behavior.

**Fig. S10.**
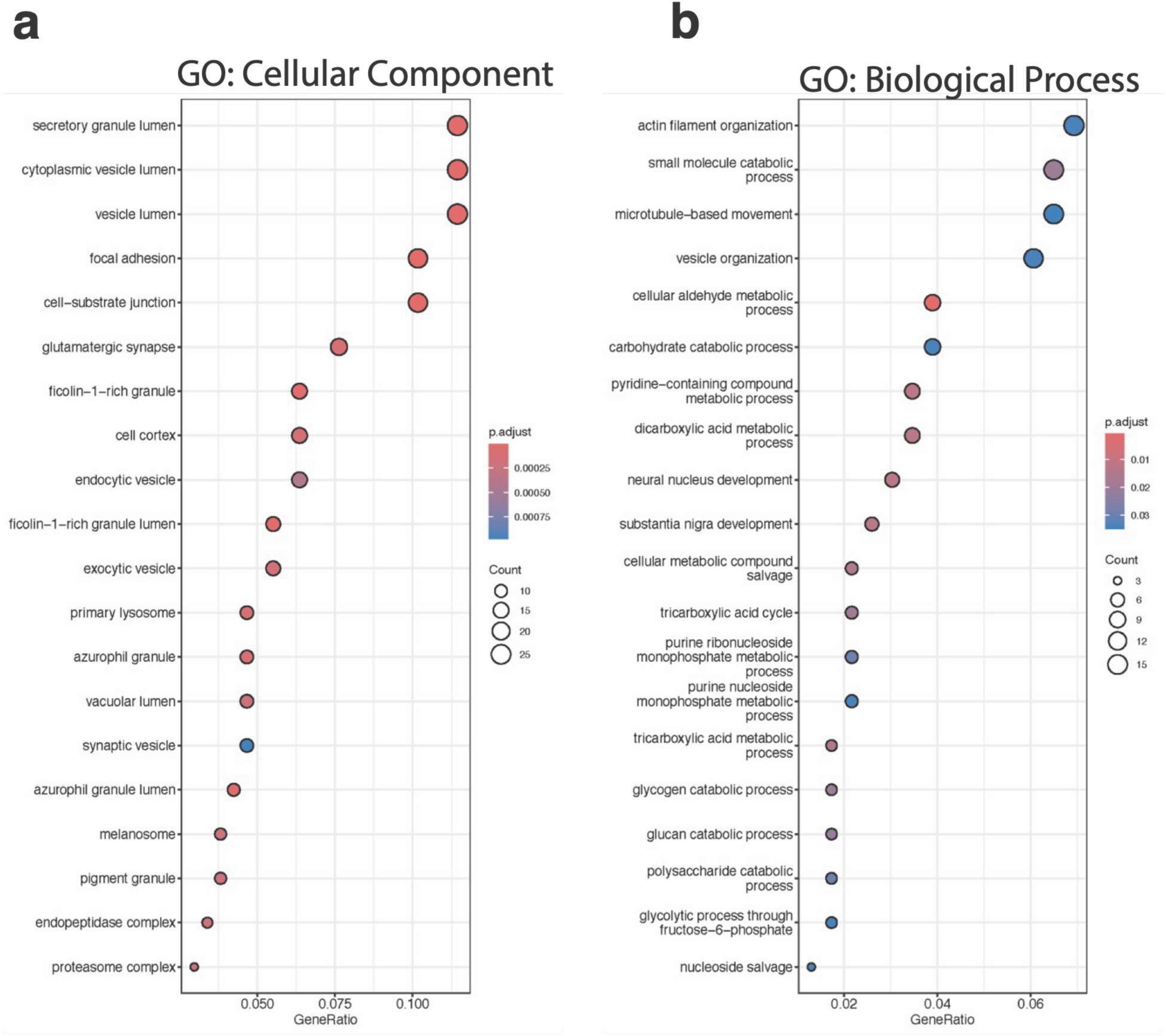
Functional enrichment analysis of proteins uniquely identified in CSF_joint_ relative to external CSF studies. Gene Ontology enrichment analysis of **(a)** cellular component and **(b)** biological process terms for the 198 proteins uniquely identified in the CSF_joint_ dataset compared with comparator CSF studies.

**Fig. S11.**
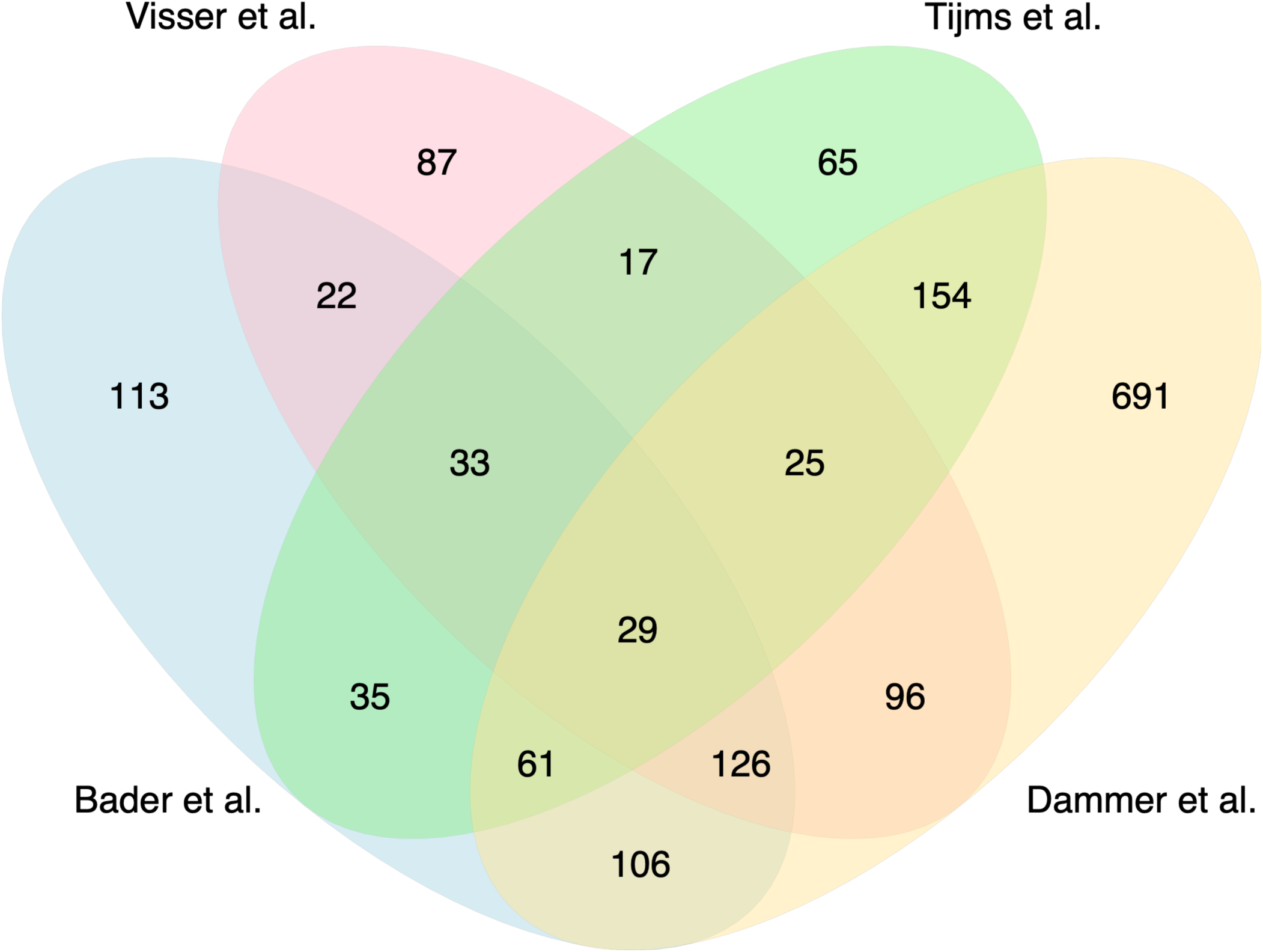
Comparison of overlap of proteins identified in external studies. Venn diagram comparing significantly differentially abundant proteins identified in four published CSF proteomic studies of AD (sample sizes ranging from 60–606 participants), illustrating the degree of overlap.

**Fig. S12.**
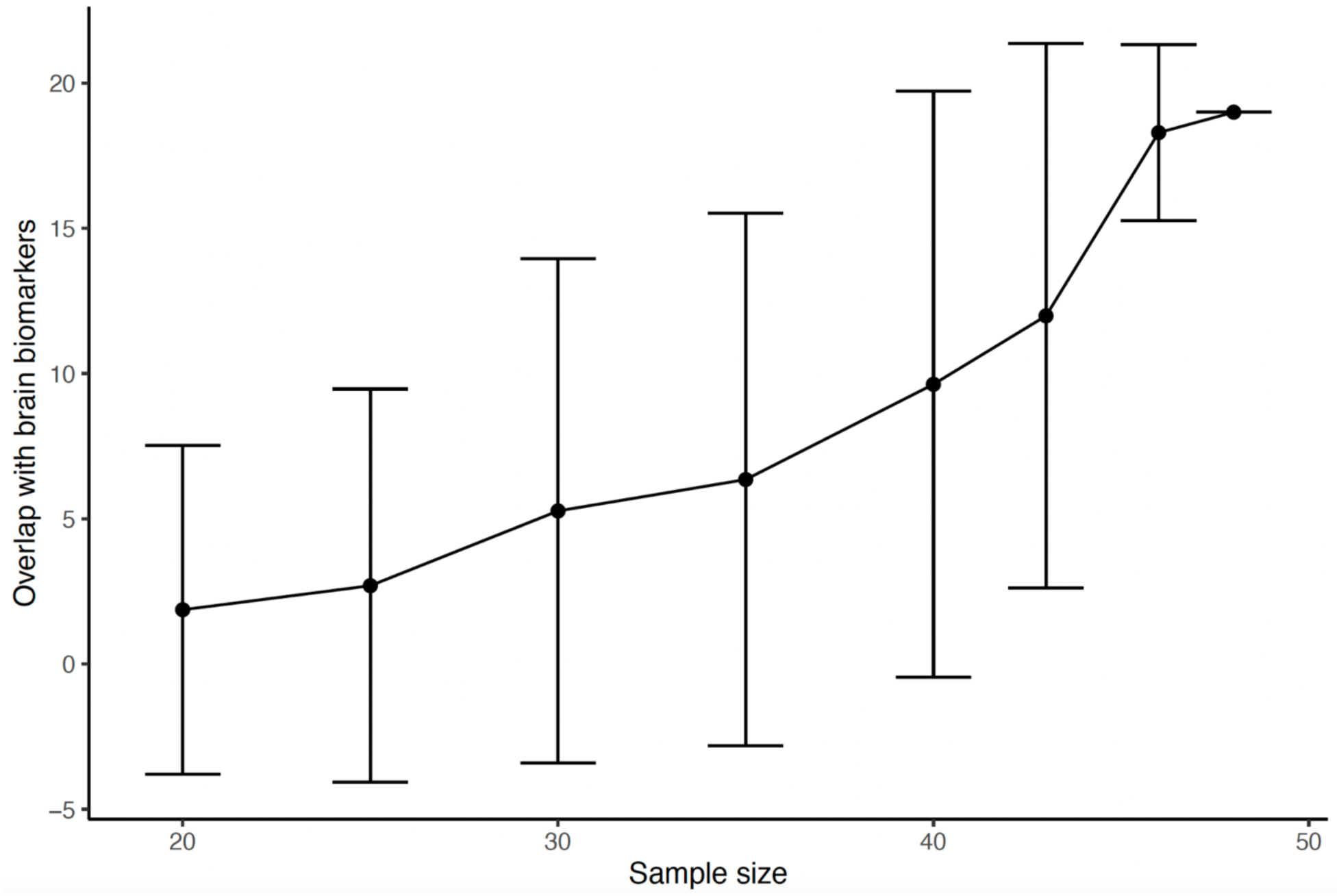
Power analysis of brain biomarker recovery by downsampling. The mean number of proteins identified in this work overlapping AD-associated brain biomarkers in Markov et al.^19^ is shown as a function of sample size. Each point represents the average overlap across 500 independent downsampling iterations, and error bars indicate ± one standard deviation.

## Methods

### Patient samples

Postmortem CSF was obtained from 48 participants in the Religious Orders Study and Memory and Aging Project (ROS/MAP): 27 with AD and 21 cognitively normal controls. Criteria used for selection: 1) advanced age at time of death (90±13 yrs). 2) a short and narrowly distributed postmortem interval (5.8±1.8 h). 3) a final consensus diagnosis (COGDX) score of either 1 (no cognitive impairment) or 4 (AD and no other cause of cognitive impairment). 4) No visual evidence of color or other obvious contaminants. By APOE genotype, 24 participants were ε3/ε3 and 24 were ε3/ε4. Among participants with AD, 12 were ε3/ε3 and 15 were ε3/ε4, whereas among controls, 12 were ε3/ε3 and 9 were ε3/ε4.

### EVps isolation and processing for LC-MS

EV_ps_ was prepared from 950 uL of the parent CSF samples using TIM4-based, MagCapture™ Exosome Isolation Kit PS Ver.2 (FUJIFILM Wako Pure Chemical Corporation, Osaka, Japan) and protocol with changes as previously described in Brown et al. (2025)^21^. CSF was clarified by centrifugation at 10,000 g for 30 min at 4-10°C. No sample produced a visible pellet. Regardless, 850 µL was transferred from the top of the centrifuge tube with care taken to avoid any disturbance of the bottom of the tube. Binding and wash steps followed the kit’s protocols. For elution, the recommended buffer was replaced with a 50% trifluoroethanol (TFE) and 0.1 M ammonium bicarbonate solution to allow direct protein extraction compatible with LC-MS analysis. Elution was performed in two rounds using the same solution: 25 µL was added in the first round and incubated on a magnetic stand and the eluate collected. This was followed by a second elution with an additional 25 µL of the same solution. The eluate was pooled, and 12 µL of 5% acetic acid was added to acidify the combined sample. Allowing for expected sample loss during processing, the recovered sample was divided into three 17.3 µL aliquots, with one aliquot used for downstream proteome analysis. Aliquots were flash-frozen in liquid nitrogen and lyophilized with all sample exposure controlled using high-purity argon introduced, as needed, via HEPA-rated gas filters. All other components of this workflow in which tubes were uncapped were conducted in a biosafety cabinet in which airborne particulates were additionally monitored in real time with an M2000 air quality meter (Temtop, San Jose, CA, USA).

### Sample workup for LC-MS

Dry EV_ps_ aliquots were resuspended in their entirety in a minimal volume of 50% TFE in 0.1 M ammonium bicarbonate with 5 mM DTT and reduced at 25°C for 1 h. Corresponding whole-CSF samples were quantified by absorbance at 280 nm, and a 50 µg aliquot of each was removed for processing and reduced with 5 mM DTT for 1.5 h at 25°C.

All samples were then alkylated with 10 mM iodoacetamide for 45 min at 25°C in the dark. Prior to digestion, EV_ps_ samples were diluted 10-fold with 0.1 M ammonium bicarbonate to reduce the TFE concentration. Modified porcine trypsin (Promega #V5113) was added at a 1:20 (w/w) enzyme:protein ratio and incubated at 37 °C overnight, after which protease activity was quenched with formic acid.

EV_ps_ peptides were dried to completion. Whole-CSF peptides were desalted by reversed-phase chromatography (Pierce peptide desalting spin columns, Thermo Fisher #89852) and then dried to completion. All samples were then resuspended in Solvent A (0.1% formic acid in Fisher Optima LC/MS-grade water), quantified by absorbance at 280 nm, and normalized to a common injection amount across samples.

### Untargeted Protein Identification and Label-free Quantification via Tandem Mass Spectrometry

Peptides were analyzed using a Thermo Scientific Ultimate 3000 RSLCnano ultra-high-performance liquid chromatography (UPLC) system coupled to a high-resolution Thermo Scientific Eclipse Tribrid Orbitrap mass spectrometer. Each sample was injected onto a nanoEase M/Z Peptide BEH C18 column (1.7 μm, 75 μm×250 mm; Waters Corporation) heated to 50°C and separated by reversed-phase UPLC at 300 nL/min: a gradient of 4–30% Solvent B (0.1% formic acid in Fisher Optima LC/MS-grade acetonitrile) over 50 min, followed by a ramp of 30–90% Solvent B over 10 min, for a 60 min total gradient. Peptides were eluted directly into the Eclipse by positive-mode nanoflow electrospray ionization with a capillary voltage of 2200 V. MS1 scans were acquired at 120,000 resolution, with an AGC target of 4e5, maximum injection time set to Auto, RF lens at 30%, and a scan range of 300–1800 m/z. Data-dependent MS2 scans were acquired in the Orbitrap at 15,000 resolution for ions exceeding a 5.0e4 intensity threshold, with AGC target set to Standard, maximum injection time set to Auto, mass range set to Normal, scan range set to Auto, isolation window set to 1.6 m/z, and a fixed cycle time of 3 s. HCD fragmentation was applied at 30% collision energy. Dynamic exclusion was set to exclude for 30 s after one observation, for charge states 2–8 only.

### Protein Identification and Quantification

Proteins were identified by database search in the FragPipe platform with the MSFragger search engine^25^, using the human proteome (Proteome ID: UP000005640) as the reference. Label-free quantification of the data-dependent acquisition (DDA) data was performed with IonQuant within the FragPipe workflow at default parameters, with match-between-runs (MBR) enabled across all samples (CSF AD, CSF Control, EV AD, and EV Control). Protein abundances were calculated as the summed intensities of the top three unique peptides per protein.

The EV_ps_ and CSF datasets were then processed separately for downstream analysis, as follows. Proteins were retained only if quantified in at least 40% of samples within a dataset. Missing values were imputed with each protein’s minimum observed intensity to represent low-abundance signal. Finally, intensities were normalized per sample to correct for systematic differences across runs.

Broadly, preprocessing of proteomic data reduces algorithmic bias toward high-intensity proteins, as absolute protein intensity is not necessarily indicative of biological relevance. CSFraw and EVraw were log2-transformed and per-protein mean-centered, removing differences in average protein intensity while retaining variation across patient samples.

### Joint and Individual Variation Explained (JIVE) analysis

To perform integrated analyses of the EV_ps_ and CSF proteomes derived from the same set of 48 individuals, we applied Joint and Individual Variation Explained (JIVE) analysis to the processed protein abundance matrices, using the r.jive package in R^23,39^. Following JIVE’s recommended preprocessing procedure, the processed data (as described above) were centered by protein to ensure comparability across datasets. To balance the contributions of the two data blocks (EV_ps_ and CSF), both matrices were scaled to have equal total Frobenius norms. JIVE was then applied to jointly decompose the EV_ps_ and CSF matrices, separating each data block into three components: (i) a joint structure capturing low-rank variation shared across data types, (ii) individual structures representing variation unique to each data type, and (iii) residual noise, with estimated ranks of 2 for the joint structure, 8 for the EV_ps_-specific structure, and 10 for the CSF-specific structure. After decomposition, each joint and individual component matrix was rescaled by multiplying back the total Frobenius norm to restore the original data scale.

### Pairwise protein-protein correlations

Pairwise protein correlations were computed as Pearson correlations across samples. For the CSF and EV data sets, correlation matrices were calculated from the preprocessed matrix used as input to JIVE. This preprocessing includes normalization and centering of the log_2_ of protein abundances to minimize the influence of global sample intensity differences and to enable direct comparison of correlation structures before and after JIVE decomposition. To compare structures across representations, proteins were hierarchically clustered on the joint-component correlation matrix using (1 − Pearson’s) correlation as the distance metric with complete linkage, and this ordering was applied to all matrices. Correlation patterns were visualized as combined heatmaps in which the upper triangle shows correlations from the joint (Fig. 2 and S5) or independent (Fig. S4) components of the JIVE decomposition and the lower triangle shows corresponding using the input data.

### Differential Abundance Analysis

Differential abundance analysis was performed separately for the EV and CSF proteomes to identify proteins showing significant differences between AD and control samples. Analyses were conducted using the original processed protein abundance matrices as well as the joint and individual component matrices derived from the JIVE decomposition, each restored to the same data scale.

For each dataset, a linear model was fitted using the limma package in R^40^ to test for group differences in protein abundance while controlling for APOE genotype, with Empirical Bayes moderation applied to improve variance estimation and statistical power. For each protein, log₂ fold changes and associated p-values were calculated. In the single-compartment analysis of original protein abundances, p-values were adjusted for multiple testing using the Benjamini–Hochberg (BH) false discovery rate (FDR) procedure^41^. In contrast, for the joint-compartment analysis based on the joint and individual component matrices, p-values were adjusted using the Benjamini–Yekutieli (BY) procedure^38^, reflecting the expected dependence structure among proteins in the low-rank space. Proteins with FDR < 0.01 were considered significantly differentially abundant.

### Functional Enrichment Analysis

Functional enrichment analysis was performed to identify biological functions and disease associations overrepresented among proteins found to be significantly differentially abundant. Gene Ontology (GO) enrichment for biological process (BP), molecular function (MF), and cellular component (CC), as well as Disease Ontology (DO) enrichment, was performed using the clusterProfiler R package^42^ with the org.Hs.eg.db annotation database and the Human Disease Ontology (HDO) as reference. Gene symbols of proteins were mapped to Entrez IDs when required. Enrichment results were adjusted for multiple testing using the Benjamini–Hochberg method, and terms with adjusted p-values < 0.05 were considered significant. Significantly enriched Gene Ontology terms were visualized using dot plots, showing the top 20 categories for each GO category. Disease Ontology enrichment results were visualized using tree plots to represent hierarchical relationships among the top 30 enriched categories.

### Permutation Analysis of JIVE

To assess the robustness of the joint structure identified by JIVE, permutation analyses were performed on the EVps and CSF protein abundance matrices. In separate analyses, the sample columns of the CSF matrix were randomly permuted while the EVps matrix was held fixed, and the sample columns of the EVps matrix were randomly permuted while the CSF matrix was held fixed. Each analysis was repeated 100 times, thereby disrupting matched CSF–EVps correspondence while preserving the internal structure of each data block. For each permuted dataset, JIVE was reapplied to determine whether joint variation depended on the true sample correspondence, and the variance explained by the joint and individual components was calculated. The distributions of explained variance across the 100 permutations for each analysis were compared with the variance explained in the original matched dataset to assess the robustness of the observed joint and individual structures.

### Leave-Two-Out Cross-Validation of JIVE

To evaluate the stability of differential abundance estimates derived from the JIVE joint component, a leave-two-out cross-validation analysis was performed by iteratively removing one AD and one control sample from the matched cohort (27 AD × 21 Control = 567 iterations). For each iteration, JIVE was re-applied to the reduced dataset to estimate the joint component, followed by differential abundance analysis using the same modeling framework. Log fold-change (log_2_FC) estimates were retained for all proteins in each iteration. The median log_2_FC across cross-validation iterations was then calculated for each protein and compared with the corresponding estimate obtained from the full dataset to assess the robustness and reproducibility of effect-size estimates across subsamples.

### Comparison to other studies

To evaluate biological relevance of biomarker candidates identified by our integrated CSF–EV framework, we compared our results with previously published mass spectrometry–based proteomic studies of Alzheimer’s disease (AD). External CSF benchmarking datasets included studies from Bader et al. (2020; n = 60)^4^, Dammer et al. (2024; n = 300)^7^, Visser et al. (2022; n = 563)^5^, and Tijms et al. (2024; n = 606)^6^, which were selected based on their large-scale CSF proteomic profiling using high-resolution mass spectrometry platforms. These studies differed substantially in cohort size, sample processing workflows, statistical methodologies, and significance thresholds (Supplementary Table S8). Because some studies included comparisons across multiple stages of cognitive decline within the AD spectrum, we focused the cross-study comparison on proteins identified as differentially abundant between AD and cognitively normal control groups (Supplementary Table S8). Differentially abundant proteins identified from our integrated CSF analysis were compared with significant proteins reported in the external studies to assess overlap with previously established AD-associated proteomic signatures. In contrast to several benchmark studies that applied relatively permissive statistical thresholds, our analysis used a stringent multiple-testing correction criterion (Benjamini-Yekutieli adjusted P value < 0.01) to define significant biomarkers. Protein overlaps were assessed at the gene-symbol level after harmonizing protein identifiers across datasets.

To further evaluate concordance with brain-derived AD proteomic signatures, we compared the identified CSF biomarkers with AD-associated proteins reported by Markov et al. (2025)^29^, which analyzed differential protein abundance in human prefrontal cortex tissue using mass spectrometry–based profiling. Protein identifiers from the brain dataset were intersected with significant proteins identified from our integrated CSF analysis and external CSF proteomics studies.

To assess the statistical power of the integrated framework for recovering established brain biomarkers, a downsampling analysis was performed across multiple cohort sizes. Samples were randomly subsampled through stratified sampling to preserve the original distribution of disease condition and genotype. Downsampled cohort sizes included n = 20, 25, 30, 35, 40, 43, 46, and 48 (Fig. S12). For each subset, differential protein expression analysis was performed using the limma framework, followed by Benjamini–Yekutieli multiple-testing correction. Proteins with adjusted P values < 0.01 were considered significant. The number of overlapping proteins between the significant CSF biomarkers and the external brain biomarker set was calculated for each iteration. This procedure was repeated 500 times independently for each sample size. Mean overlap and standard deviation across iterations were computed to estimate the relationship between sample size and the ability to recover previously reported brain-associated AD biomarkers (Fig. S12).

## Acknowledgements

This work was supported by a Warren Alpert Foundation grant awarded to GG, and the National Institute on Aging grant R01AG068285 awarded to ADM. Special thanks to Jeremy Balsbaugh and Jen Liddle at the University Connecticut Proteomics & Metabolomics Facility, supported in part by NIH High-End Instrumentation Award 1S10-OD028445.

## Author Contributions

- Conceptualization: BRPB, XL, ADM, GG
- Methodology BRPB, XL, ADM, GG, MG
- Investigation BRPB, ADM, MG
- Writing, original draft: BRPB, XL, ADM, GG
- Writing, review and editing: BRPB, XL, ADM, GG
- Funding acquisition: ADM, GG
- Supervision: ADM, GG

## Data and code availability

- Data for this study are publicly available as of the date of publication. The mass spectrometry proteomics data will be deposited to the Synapse.
- All original code for data processing and performing downstream statistics can be found at https://github.com/G2Lab/Alzheimers_CSF_EV_Proteomics_Integration
- Any additional information required to reanalyze the data reported in this paper is available from the lead contact upon request.

## Notes

https://github.com/G2Lab/Alzheimers_CSF_EV_Proteomics_Integration

### Competing Interest Statement

The authors have declared no competing interest.

